# Architectural experience: clarifying its central components and their relation to core affect with a set of first-person-view videos

**DOI:** 10.1101/2022.04.05.487021

**Authors:** Lara Gregorians, Pablo Fernández Velasco, Fiona Zisch, Hugo J. Spiers

## Abstract

When studying architectural experience in the lab, it is of paramount importance to use a proxy as close to real-world experience as possible. Whilst still images visually describe real spaces, and virtual reality allows for dynamic movement, each medium lacks the alternative attribute. To merge these benefits, we created and validated a novel dataset of valenced videos of first-person-view travel through built environments. This dataset was then used to clarify the relationship of core affect (valence and arousal) and architectural experience. Specifically, we verified the relationship between valence and fascination, coherence, and hominess - three key psychological dimensions of architectural experience which have previously been shown to explain aesthetic ratings of built environments. We also found that arousal is only significantly correlated with fascination, and that both are embedded in a relationship with spatial complexity and unusualness. These results help to clarify the nature of fascination, and to distinguish it from coherence and hominess when it comes to core affect. Moreover, these results demonstrate the utility of a video dataset of affect-laden spaces for understanding architectural experience.

**Highlights:**

- Developed a video database of first-person-view journeys through built environments
- We explored how core affect and architectural experience relate through the videos
- Previous results are supported: valence ties to fascination, coherence and hominess
- Arousal correlates only with fascination, and not coherence or hominess
- Arousal and fascination are tied to spatial complexity and unusualness

## 1. Introduction

Architectural experience is at once commonplace and unique. Commonplace, because of the large amount of time that humans spend in architecture (defined here as space inside and between buildings; Evans, 2000); unique, because the experience of architecture is unlike other aesthetic experiences. One does not just contemplate a building the way one might contemplate a painting. One goes through a building, explores a building, dwells in a building, and eventually leaves it. Architecture is multisensory and dynamic, generating atmosphere which surpasses just the visual or aesthetic, and hinges on human-environment interactions (see texts on architectural phenomenology e.g. Böhme, 1993, 2018; Zumthor, 2006; Robinson, 2011, 2021; Pallasmaa, 2012, 2014). Essential to our experiences of buildings are the paths we take and the views our gaze captures.

However, what makes architectural experience unique is also what makes it hard to study. While paintings or musical pieces provide suitable experimental stimuli, no easy substitute exists for architecture, where space itself ‘determines the aesthetic value of the building’ (Van Doesburg, 1919). Empirical researchers can either use still images of buildings, which do not transmit, for example, the dynamic nature of walking through buildings, or they can use virtual reality environments, which in their distinctive computer-generated texture engender a different atmosphere from being in physical architectural spaces (Böhme 1993, 2016, 2018; Mattens, 2011). In the face of this challenge, we have developed and tested a dataset of videos of trajectories through real-life built environments which can be used for empirical research on the experience of built environments. This provides a direct step-up from images by incorporating dynamic movement, but is still as easily accessible as image databases, and useable for online testing and functional magnetic resonance imaging (fMRI).

Here, we introduce this dataset and examine the nature of architectural experience, which relates to the experience of encountering a built environment with a focus on the space itself – for example, exploring a new building to see what is there. Encompassed within this paper’s analysis of architectural experience are aesthetic valuation, affective response and spatial mapping – on entering a space we take in sensory information; evaluate the space’s characteristics, beauty and how it makes us feel; but we must also comprehend the space to fulfil the functional goal of moving through it (Zisch, Gage, Spiers, 2014; Coburn, Vartanian & Chatterjee, 2017; Gregorians & Spiers, in press). Whilst aesthetic and affective experience are often considered in tandem, spatial cognition tends to be considered quite separately. Nonetheless, all three processes are embedded in real-world spatial experiences, affecting our ability and desire to dwell, explore or navigate. As Pessoa (2008, 2013) argues, emotion and cognition are highly integrated in many functions including visual processing. In this study, we evaluate the interactions of valence and arousal (the two dimensions of core affect; Russell, 2003) with fascination, coherence and hominess (three central components underlying aesthetic responses to architectural experiences; Coburn et al., 2020), and also two factors we propose may be important: spatial complexity (how complex the layout is) and unusualness. We employ our new dataset of videos to further our understanding of architectural experience and its affective underpinnings. Furthermore, we show how this dataset was analysed to derive a subset of videos that are distinctly valenced, providing a streamlined video database that can be used for future studies exploring affective responses to the built environment.

### 1.1. Empirical aesthetics, architectural experience and emotion

Aesthetic experience hinges on the perception and evaluation of objects – objects that may not pose value to survival or function yet can hold high or low value in beauty, harmony, awe (etc.) – and involves an interaction of sensory, emotional and personal reactions (Jacobsen et al., 2004; Jacobsen, 2006; Vessel, Starr & Rubin, 2012, 2013). Architectural experience innately involves aesthetic processing, and emotion has been identified as a key pillar in both (Chatterjee & Vartanian, 2014; Coburn, Vartanian, & Chatterjee, 2017), influencing the production of presence and atmosphere (Griffero 2016), along with meaning-making (Gumbrecht 2004) and ties to sensorimotor systems (Chatterjee & Vartanian, 2014). Nadal and colleagues (2008) argue that ‘what sets aesthetic preference apart from other cognitive processing of visual stimuli is precisely the engagement and interplay of additional non-perceptual processes, such as emotions.’ Similarly, the feeling of pleasure is considered one of the most important facets of aesthetic experience (Brielmann & Pelli, 2017). Crucially for the built environment context, not only are emotions central to the valuation of the aesthetic object, but they also drive behaviours — attracting or repelling us, both physically and metaphorically (Yazdani et al., 2012). Thus, aesthetic emotions play a key role influencing decision making, including how we choose to enter and move through spaces (Brown et al., 2011).

Much of the recent scientific advances regarding architectural experience can be framed within the broader approach of ‘neuroarchitecture’ (see e.g. Eberhard 2007, 2009; Mallgrave 2010, 2013; Metzger, 2018) and ‘neuroaesthetics’ (Coburn, Vartanian, & Chatterjee, 2017; Robinson & Pallasmaa, 2015). Neuroaesthetics tries to clarify the neural substrata of aesthetic responses. Research has gathered that aesthetic experience involves three key neural systems of sensation/perception, understanding and reward (Chatterjee & Vartanian, 2014; Belfi et al., 2019), and that the ‘core circuit for aesthetic processing’ involves the orbitofrontal cortex, basal ganglia, anterior insula, and the rostral part of the cingulate cortex (Brown et al. 2011). Research using functional magnetic resonance imaging (fMRI) has found that an ‘evaluative’ network of frontal regions including nodes of the default mode network (DMN) – thought to support inward contemplation – tap into the personal nature of aesthetic experience, showing a step-like pattern of activation that signals when stimuli is not just liked but found moving (Vessel, Starr & Rubin, 2012). Applied to architecture, fMRI has helped identify patterns of neural activity associated with specific architectural styles (Choo, Nasar, Nikrahei, & Walther, 2017) and distinctive neural responses to built environments with curvilinear contours (Vartanian et al., 2013). The desire to leave enclosed rooms, compared to open rooms, has been found to activate the anterior midcingulate cortex (Vartanian et al., 2015), and broader DMN engagement linked to aesthetic appeal for artwork has also been found to extend to images of architecture (Vessel et al., 2019).

However, a limitation with past fMRI research examining architecture is that there is somewhat of a lacuna when it comes to the experimental psychology of architecture (Graham, Gosling, & Travis, 2015). Architects and philosophers have long theorised about the nature of our experience and aesthetic appreciation of architecture (Vischer 1893; Lipps 1897; Bachelard 1958, Burke 1958, Pask 1971; Scruton 1980, Alexander 2002; Semper 2004; Hill 2006; Schützeichel, 2013). Indeed, Jacobsen (2006) developed a framework outlining how the ‘psychology of aesthetics’ can be approached from seven perspectives, uniting to create a more holistic understanding of aesthetic processing that considers ‘stimulus-, personality- and situation-related factors’. However, empirical aesthetics lacks the necessary range of behavioural experiments required to construct psychological theories that can then be applied to neuroscience (Coburn, Vartanian, & Chatterjee, 2017).

A notable recent advance comes from a study by Coburn and colleagues (2020). Participants rated images of built environments on 16 aesthetic dimensions of architectural experience (e.g. beauty, relaxation, uplift…). A Psychometric Network Analysis (PNA) and Principal Components Analysis (PCA) of the ratings identified three dimensions that explained 90% of the variance in ratings: coherence, fascination and hominess. Coherence corresponds to the ease with which a participant comprehends an environment, fascination is the interest that the environment generates, and hominess is the degree to which the environment feels like a personal space. The authors then reanalysed fMRI data from two previous studies (Vartanian et al., 2013, 2015) where participants had given beauty or approach-avoidance judgments on the same images of built environments, to now explore neural activation in relation to coherence, fascination and hominess. Results showed that fascination covaried with activation in the right lingual gyrus, that coherence covaried with activation in the left inferior occipital gyrus (but only when participants were asked to judge beauty), and that hominess covaried with activation in the left cuneus (but only when participants were asked to make approach-avoidance decisions). These results suggest that distinct brain regions support the processing of these three psychological dimensions, supporting the view that coherence, fascination and hominess are distinct constructs central to architectural experience.

### 1.2. Spatial cognition, architectural experience and emotion

The built environment is also centrally important to the field of spatial cognition; from this perspective, we know a great deal about how the brain maps spaces to allow successful navigation (Nyberg, Duvelle, Barry & Spiers, 2022). Designed features of the environment are known to be central to the ease and success of this, e.g. wall boundaries help build mental representations of a space (Jeffery, 2021). The processing of these features is embedded in early perceptual processing, with perception, cognition and action unfolding jointly in respect to the changing body-environment relationship (Djebbara et al., 2019, 2021). However, studied environments in most experiments are often considered as more abstract spaces, rather than as specific and contextualised spaces (Epstein, Patai, Julian & Spiers, 2017). In essence, the importance of architectural detail and affective experience have largely been neglected within spatial cognition.

However, there have been some suggestions as to how spatial processing and affective responses intertwine. For example, environmental features that are key to navigation (e.g. route choices, novelty of environments) can provoke physiological stress responses in rodents (Bailey et al., 2006; cited in Sternberg & Wilson, 2006). In humans, the ‘affective charge’ of landmarks has been shown to be important for wayfinding: emotionally-laden landmarks are remembered better (Balaban, Roser and Hamburger 2014), wayfinding performance and recognition improve with negatively-laden landmarks (Balaban et al 2017), and positively-laden and high arousing landmarks improve the drawing of routes (Ruotolo, Claessen, van der Ham 2019; Ruotolo, Sbordone, van der Ham, 2021). Palmiero and Piccardi (2017) found emotion-laden landmarks to enhance egocentric-based topographical memory, while positive-laden landmarks specifically enhance allocentric-based topographical memory. In studies on London taxi drivers, emotion was identified as a key thought category when recalling journeys through virtual London (Spiers and Maguire 2008). There appears to be evidence for the emotional nature of human wayfinding, as well as the existence of specifically spatial affective states, such as disorientation, agoraphobia or the aha! experience that comes with sudden reorientation (Sternberg & Wilson, 2006; Jeffery, 2019; Fernández Velasco and Casati, 2020; Montello, 2020; Charalambous, Hanna, Penn, 2021). What such studies so far have neglected is the potential impact of both affect and arousal. Some experiences may be negative or positive, but have little arousal associated with them, while others may drive a heightened state of arousal (e.g. Russell et al., 1989). Understanding how affect and arousal are related to architectural experiences is an important, as yet unexplored, question.

While we focus here on theory and methods emerging from empirical aesthetics, it is important to emphasise that architectural experience should also be understood in the broader context of spatial cognition as well as affective processes. As such, this study considers architectural experience in relation to core affect and parameters pertinent to aesthetic evaluation and spatial mapping.

### 1.3. Aims and hypotheses

There are four interrelated aims in this study.

#### 1. To develop a dataset of valenced videos of trajectories through real-life built environments that can be used to study architectural experience

The completion of the first aim is outlined in the methods section, and its outcome serves as the base of the following two aims.

#### 2. To use this dataset to understand the relationship between affect and architectural experience

An online survey is used to investigate how the two core dimensions of affect (valence and arousal) relate to other aspects of architectural experience, asking specifically:

A. Is valence correlated to fascination, coherence and hominess?
B. Is arousal correlated to fascination, coherence and hominess?
C. Do aesthetic and affective dimensions correlate with spatial complexity and unusualness?
D. How are valence and beauty related?

The study by Coburn et al. (2020) strongly predicts that the answer to (A) will be positive. Finding this to be the case with our novel dataset would, on the one hand, further support the importance of these three dimensions of architectural experience, and, on the other, support the effectiveness of our dataset in empirical research, as it successfully reproduces clear findings in previous work. As for (B), the outcome is more speculative. We reason that if affect is of chief importance to architectural experience, and that fascination, coherence and hominess are all related to valence, then arousal (the other dimension of core affect) might help set these three dimensions apart and further clarify the nature of architectural experience. For (C), we speculate that videos that are more spatially complex and unusual will be considered more fascinating and arousing. This is predicated on the basis that unusual environments should trigger a novelty response which would be linked to arousal (Mendes et al., 2007), and that arousal (both self-induced and landmark-laden) can alter spatial judgement, planning and memory (Gardony et al., 2011; Brown et al., 2020; Ruotolo, Sbordone, van der Ham, 2021). Moreover, unusualness (or novelty), complexity and arousal are found to relate or influence architectural engagement (Berlyne, 1970; Berlyne, 1973; Oostendorp & Berlyne, 1978; Xylakis, Liapis & Yannakakis, 2021). We also speculate that environments that are more complex in their layout may tend to be more fascinating compared to very simple environments given that Coburn et al. (2020) find spatial complexity to tie heavily to fascination. Furthermore for (D), we explore the relationship between beauty and valence to consider how responses to these two interrelated but different components of aesthetic experience relate to one another, and whether this has implications on how the video dataset can be used for study. We hypothesise that beauty and pleasure ratings will be highly correlated, as has been shown across different stimuli sets (e.g. Vartanian et al., 2013; Brielmann & Pelli, 2017; Brielmann & Pelli, 2019).

Table 1 outlines correlations that are predicted for questions (A-C) based on literature and previous findings, and those that are speculated.

**Table 1:** Predicted Pattern of Correlations

|  | Valence | Fascination | Coherence | Hominess | Spatial comp. | Arousal | Unusualness |
| --- | --- | --- | --- | --- | --- | --- | --- |
| **Valence** |  |  |  |  |  |  |  |
| **Fascination** | Rep. |  |  |  |  |  |  |
| **Coherence** | Rep. | Rep. |  |  |  |  |  |
| **Hominess** | Rep. | Rep. | Rep. |  |  |  |  |
| **Spatial comp.** |  | Pred. |  |  |  |  |  |
| **Arousal** |  | Pred. |  |  | Pred. |  |  |
| **Unusualness** |  | Pred. |  |  |  | Pred. |  |

All correlations are predicted to be positive. Spatial comp. = spatial complexity, Rep. = correlations we are seeking to replicate, $p < .05$ set for significance. Pred. = predicted correlations, set at $p < .001$ with Bonferroni correction (see 1.3. Aims and hypotheses). Other cells in the matrix are not predicted and significance threshold set at $p < .001$ with Bonferroni correction. Not shown, we examined the correlation between the presence of people and the presence of blue/green space, $p < .05$ . This was not a hypothesis, but an assessment of the status of the video content for consideration towards confounds in the ratings.

#### 3. To use this dataset to examine whether architectural experience is modulated by self-rated navigational abilities

There is wide variation in navigation ability which impacts people’s ability to make sense of a space and navigate (e.g. Weisberg & Newcombe, 2018; Coutrot et al., 2022; Spiers et al., 2021; Hegarty et al., 2022; Ishikawa, 2022), and their use of cues when learning environments (Ishikawa & Nakamura, 2012). We explored whether self-assessed navigational ability influences responses to architectural environments. We hypothesise that: a) more confident navigators will have more positive responses to complex spaces that they might find easier to make sense of, and that b) poor navigators may find spatial complexity more arousing due to the challenge of making sense of the space. These hypotheses are based on assumptions about how spatial abilities might impact the appreciation of space rather than evidence from past research. Thus these predictions are more speculative.

#### 4. To identify a subset of affectively charged videos that vary in valence but that elicit substantially consistent valence ratings across participants, which can be used as stimuli for future research

Finally, we identify which videos from the dataset can be deduced as distinctly positive or negative in valence, and can therefore be part of a streamlined database of stimuli that can be used for future research. We believe that a verified database of valenced video stimuli will be useful for exploring the dynamic experience of architectural space.

## 2. Methods

### 2.1. Building the dataset

There were seven steps in the development of the video database:

*Stage 1)* We searched for videos that filmed first-person-view trajectories through built environments, initially sourcing ∼200 videos from YouTube.

*Stage 2)* Two pairs of coders from the research team evaluated whether the videos were negatively or positively valenced on a scale from −10 to 10. We also recorded the presence of people and green or blue space in the videos. We discarded videos if there was no agreement between the two coders with regards to valence (firstly if scores differed by more than 5, secondly over qualitative discussion), and videos that were deemed problematic in other respects (e.g. distorted lens).

*Stage 3)* We selected and edited videos to produce clips 31-40s long. Figure 1 shows a storyboard of stills from one video to outline an example journey

**Figure 1:**
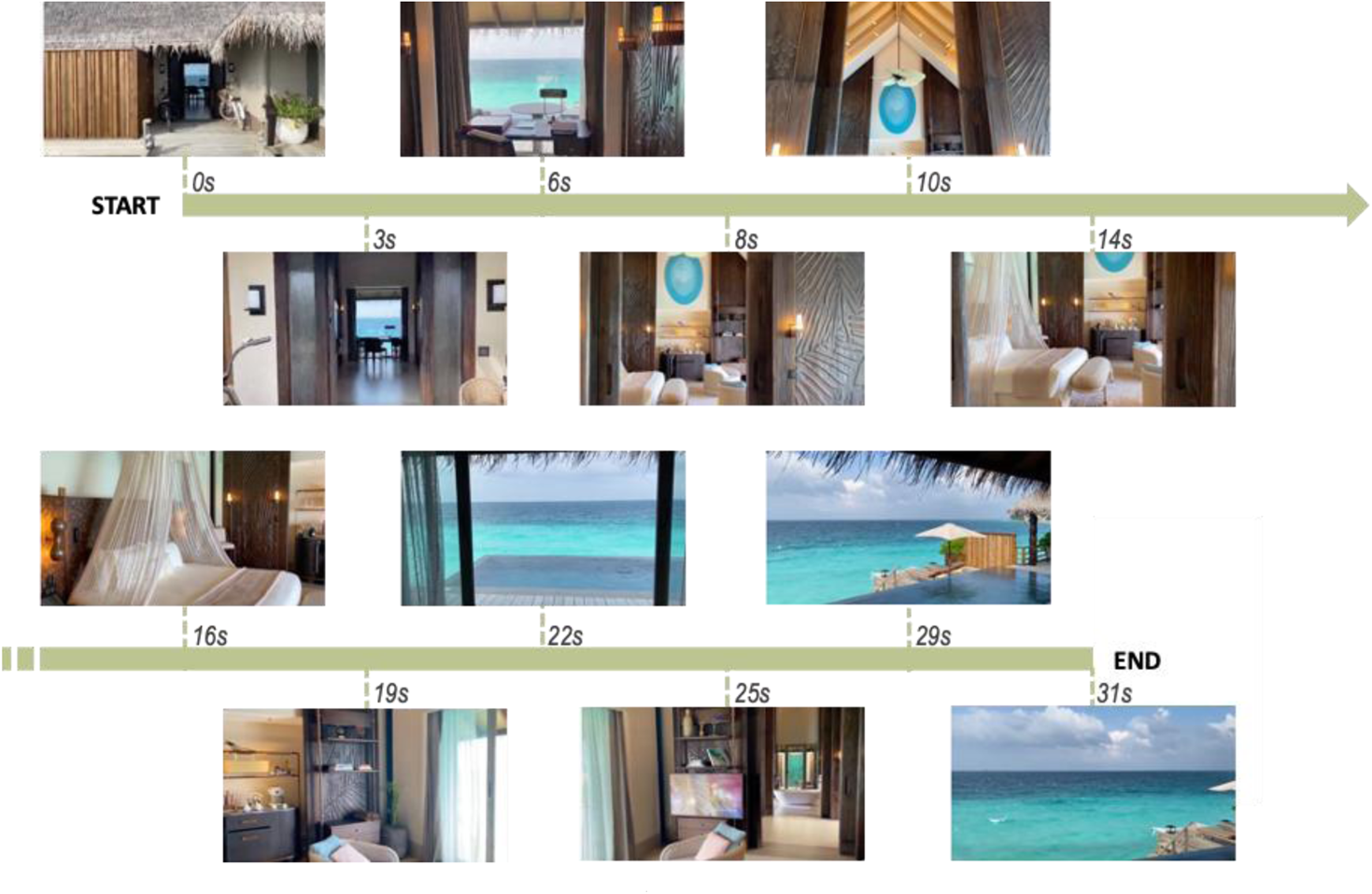
Storyboard showing the timeline of a video.

*Stage 4)* Each of the resulting clips were assessed by an architectural expert to identify if they contained architectural events of interest (e.g. changes in scale, confinement, connectivity, lines of sight or depth of view). We adopted this method following studies that have used analysis by experts to evaluate specific qualitative criteria from stimuli (e.g. Antony et al., 2021; Erkan, 2018). We discarded videos that had no architectural events of interest. This allows for our dataset to be rich with architectural information that can be analysed in future research, to better understand responses to specific architectural qualities. This also follows previous work (e.g. Coburn et al., 2020) where stimuli buildings were diverse in architectural properties.

*Stage 5)* We categorised the building types filmed in the clips to ensure that our set included a broad variety of spaces. A challenge for this is that, to our knowledge, there is no systematic and detailed taxonomy of building types. Tversky and Hemenway (1983) argue that the best categorization can be found in architectural resources (e.g. textbooks). Consequently, we conducted an overview of such resources (e.g. de Chiara and Callender, 1973) to construct a categorization of built environments (e.g. health care, commercial, educational, residential, etc.). Our search found 42 different building types, and the final database at the end of the process covered 30 building types.

*Stage 6)* We eliminated some clips to ensure that our set included an even distribution of positively and negatively valenced videos (as deduced by the research team in stage 2), and within the positive and negative subsets, that there be an even distribution of videos with people, and videos with green or blue space to control for these factors. As the purpose of this study was to examine architectural experience, we sought to minimise the effect of green or blue space, which is known to be able to impact user experience (e.g. Joye, 2007; Reeve et al., 2012). Therefore, our final selection of clips purposefully included videos with good, bad and no green or blue space across both (hypothesised) positive and negatively valenced spaces. We found no impact of green/blue space or people on key findings. The overall process resulted in a final set of 72 videos (table 2).

**Table 2:** Distribution of stimuli set accounting for valence, green/blue space and presence of people

|  |  | Hypothesised valence |  |
| --- | --- | --- | --- |
|  |  | Positive | Negative |
| Attributes | Green/ blue space | 13 | 13 |
|  | With people | 22 | 22 |
|  | **Total** | 36 | 36 |

*Stage 7)* The 72 clips were broken into 4 blocks of 18 videos. Blocks were designed to equally distribute as closely as possible clips that were positive, negative, with green/blue space, and with people. The spatial layout complexity of each space was also evaluated by an architectural expert in order to distribute simple and complex spaces between the blocks. Within blocks, the order of videos was set so that videos alternated between positive and negative valence randomly, with no more than two clips of the same valence following one another. Within block orders remained fixed across participants.

### 2.2. Participants

Power analysis using G*Power 3.1.94 indicates a sample size of 60-182 is needed to run two-tailed correlations with effect size = 0.3-0.5, power = 0.8 and *ɑ* = .001. We recruited 78 participants to take part in an online survey (Survey 1) via Prolific and the SONA platform of the first author’s host institution. Participants were automatically assigned to one of four groups, indicating which video blocks they would be rating. Each group had a different order of video blocks (labelled blocks A-D) for partial counterbalancing. To ensure that the survey completion time was under 1 hour and to prevent overstimulation, each participant group saw two blocks of the possible four blocks of videos (e.g. participants in Group 1 watched block A and then block B, and participants in group 3 watched block C and then block D).

Participants indicated whether they were architectural experts or students, as research indicates that architectural experts view spaces differently to the average person, with neural activity in aesthetic judgement and reward processing differing to that of a non-expert. Therefore we excluded responses from this group (Kirk et al., 2009; Vartanian et al., 2019; Wiesmann & Ishai, 2011). One participant identified as an architectural expert or student, and so their ratings were removed from the set. Participants also completed the survey-based strategy questions from the Navigational Strategy Questionnaire (NSQ), to create a mean NSQ score per participant which is used to benchmark their spatial skills (Zhong, 2013; Zhong & Kozhevnikov, 2016; Zhong, 2020). NSQ data shows mean scores were similar across groups. Full demographic information can be found in table 3. Of the 78 participants, one was excluded due to their architectural expertise, and four were excluded for not completing the task sufficiently, resulting in data from 73 participants.

**Table 3:** Demographics overview for 73 included participants of Survey 1

| Age |  | Gender |  | Origin |  | No. participants |  | NSQ score |  |  |
| --- | --- | --- | --- | --- | --- | --- | --- | --- | --- | --- |
| 18-25 | 61 | Male | 16 | UK | 30 | Group A | 19 | Group A | Max | 3.55 |
| 25-35 | 6 | Female | 57 | East Asia | 24 | Group B | 20 |  | Min | 1.45 |
| 35-45 | 3 |  |  | Europe | 10 | Group C | 20 | Group B | Max | 4.45 |
| 45-55 | 1 |  |  | Southeast Asia | 4 | Group D | 14 |  | Min | 2.09 |
| 55+ | 2 |  |  | South Asia | 2 |  |  | Group C | Max | 3.73 |
|  |  |  |  | North America | 2 |  |  |  | Min | 1.27 |
|  |  |  |  | Middle East | 1 |  |  | Group D | Max | 3.36 |
|  |  |  |  |  |  |  |  |  | Min | 1.64 |

This process was then repeated to recruit a further 16 participants, who would partake in a second survey (Survey 2), where valence was replaced with a beauty rating. A smaller sample size was needed for this survey, as its results are solely used to compare beauty and valence ratings across samples, rather than all seven qualities included in the ratings. No participants were architectural experts. Full demographic information can be found in table 4. Ethics approval was given by the first author’s institution prior to either survey’s dissemination.

**Table 4:** Demographics overview for 16 participants of survey 2

| Age |  | Gender |  | Origin |  | No. participants |  | NSQ score |  |  |
| --- | --- | --- | --- | --- | --- | --- | --- | --- | --- | --- |
| 18-25 | 16 | Male | 2 | UK | 2 | Group A | 4 | Group A | Max | 3.73 |
|  |  | Female | 14 | East Asia | 6 | Group B | 4 |  | Min | 2.09 |
|  |  |  |  | Europe | 4 | Group C | 4 | Group B | Max | 4 |
|  |  |  |  | Southeast Asia | 1 | Group D | 4 |  | Min | 3.27 |
|  |  |  |  | South Asia | 3 |  |  | Group C | Max | 3.55 |
|  |  |  |  |  |  |  |  |  | Min | 2.27 |
|  |  |  |  |  |  |  |  | Group D | Max | 4.36 |
|  |  |  |  |  |  |  |  |  | Min | 1.82 |

Participants were provided with an information sheet and they signed consent. They then proceeded to answer the pre-test questions outlined in 2.2. In Survey 1 (N=73), participants were asked to rate the built environments filmed in the videos for the following qualities: valence, arousal, spatial layout complexity, fascination, coherence, hominess, and unusualness. Valence and arousal reflect core affect; fascination, coherence and hominess speak to the relevant work of Coburn et al. (2020); and spatial complexity and unusualness are included to consider factors related to spatial cognition and memory. In Survey 2 (N=16), the rating on valence was replaced with a rating on beauty. Participants received training on how to use the rating scales, and reference images were provided for the seven qualities (see Appendix). Participants were informed that these reference images did not substitute as definitions for the terms, but purely served as useful visual descriptors. These images were important as piloting suggested that some of the less commonplace terms (e.g. coherence) could be better understood through visual aids. Images also helped distinguish between otherwise confusable terms (e.g. spatial complexity and visual complexity).

After completing training, ratings were collected. A fixation cross would appear on screen, followed by the first clip, and then participants would be asked to provide the seven ratings for that clip. This process repeated for each video in their two assigned video blocks. This survey procedure was finalised after an initial stage of piloting and modification; the final procedure for Survey 1 is outlined in figure 2.

**Figure 2:**
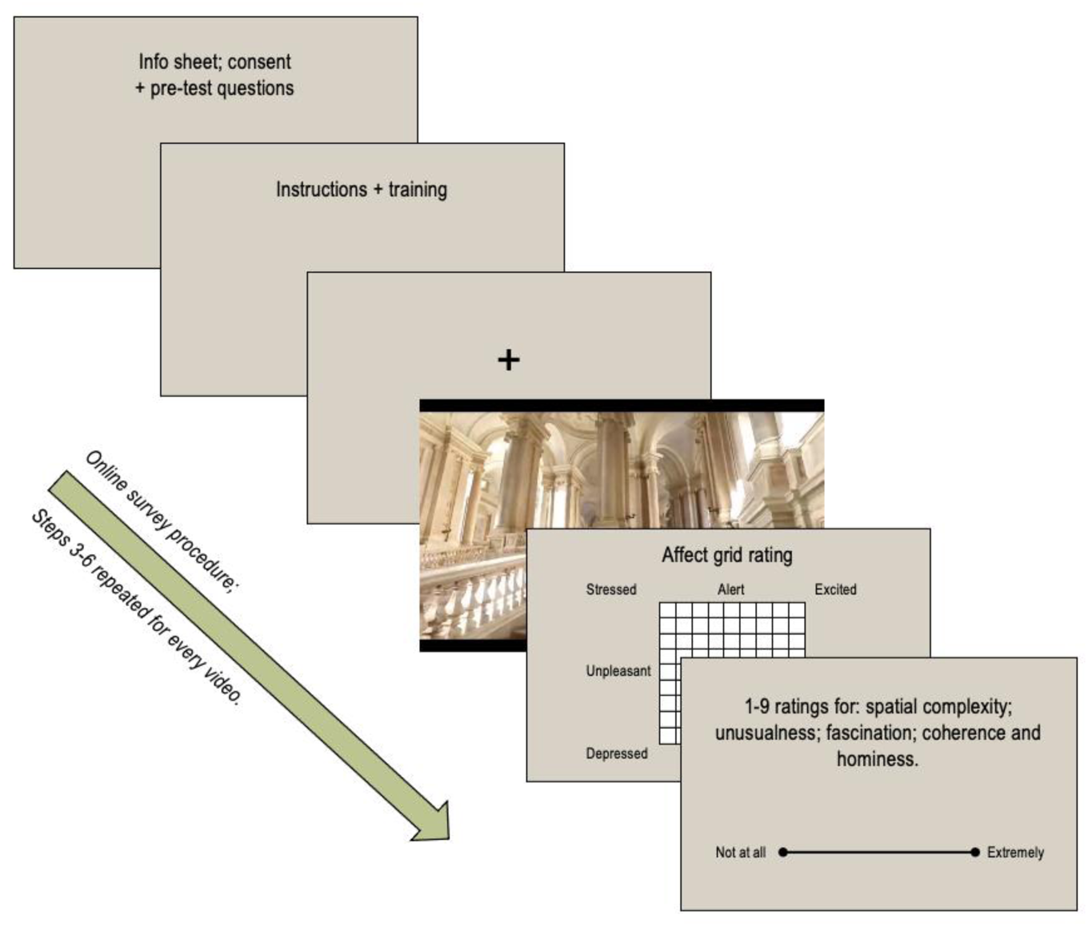
Overview of Survey 1 procedure, through which a subset of 63 videos were selected as distinctly valenced. All steps were self-paced, except fixation cross (250ms) and movies (varied 31-40s) which could not be paused or skipped.

Ratings for valence and arousal were given on the 9×9 affect grid created by Russell and colleagues (1989; see figure 3). Ratings for spatial layout complexity, fascination, coherence, hominess and unusualness were given on a discrete 1-9 scale (“not at all – extremely”), to replicate the grid’s 1-9 scale length. The affect grid has been verified against other affect-rating scales, and has been previously employed to capture affective responses to visual stimuli (Killgore, 1998; Gerber et al., 2008; Sato et al., 2020). The grid provides an understanding of valence and arousal as a 2-dimensional model consisting of two independent but related factors. In the model, valence corresponds to the degree of pleasantness-unpleasantness that results from the stimuli, and arousal corresponds to the state of inner activation or alertness. In Survey 2, the affect grid was not used, but rather, separate 1-9 scales were used to capture beauty and arousal ratings (in addition to the other five ratings), as the affect grid was not designed to assess this dimension of aesthetic experience.

**Figure 3:**
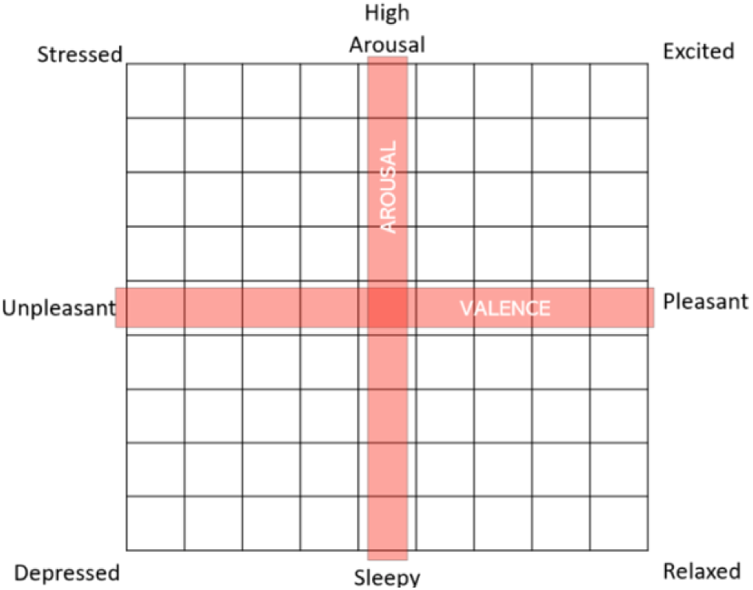
Russell et al. (1989) affect grid, modified for study (central arousal and valence axes labels included only in training phase for descriptive purposes).

## 3. Analysis and results

To explore the relationship between core affect and architectural qualities, we will focus on the results of Survey 1, where 73 participants rated 36 videos each, resulting in a total of 2625 data points (three data points were missing due to technical error). Data across all videos and participants were combined, and data analysis was carried out at the item level by calculating the average rating for each video on all seven rating scales. Firstly, we looked at which videos scored highest and lowest on average in each of the seven categories. Then we explored the relationships between these seven categories across all 72 videos by producing a correlation matrix. Several of these relationships were predicted (see table 1). Correlations which were replications were considered at the threshold p<.05, whilst all other correlation analyses were Bonferroni corrected for multiple comparisons, p<.001 (36 comparisons). Next, we compared results from Survey 1 and Survey 2 to deduce whether beauty and valence ratings were similar; if a place is rated highly positive, is it likely to be rated highly beautiful? Results from Survey 1 were then reanalysed to compare ratings given only by high and low NSQ scorers, to consider whether person-specific criteria may impact responses. As exploratory analysis, correlations were again Bonferroni corrected for multiple comparisons, p<.001. We then analysed the effects of green/blue space and the presence of people to check if these features were substantially influencing ratings. Finally, as one of the aims of this paper is to generate a dataset of valenced videos, we ran a two-tiered verification process for each video, to curate a subset of videos that were distinctly valenced.

### 3.1. Highest and lowest ratings

Across all videos in Survey 1, mean ratings for each of the seven qualities for each video were calculated, to identify which spaces scored highest or lowest. Figure 4 gives representative still images and descriptions for these respective spaces.

**Figure 4:**
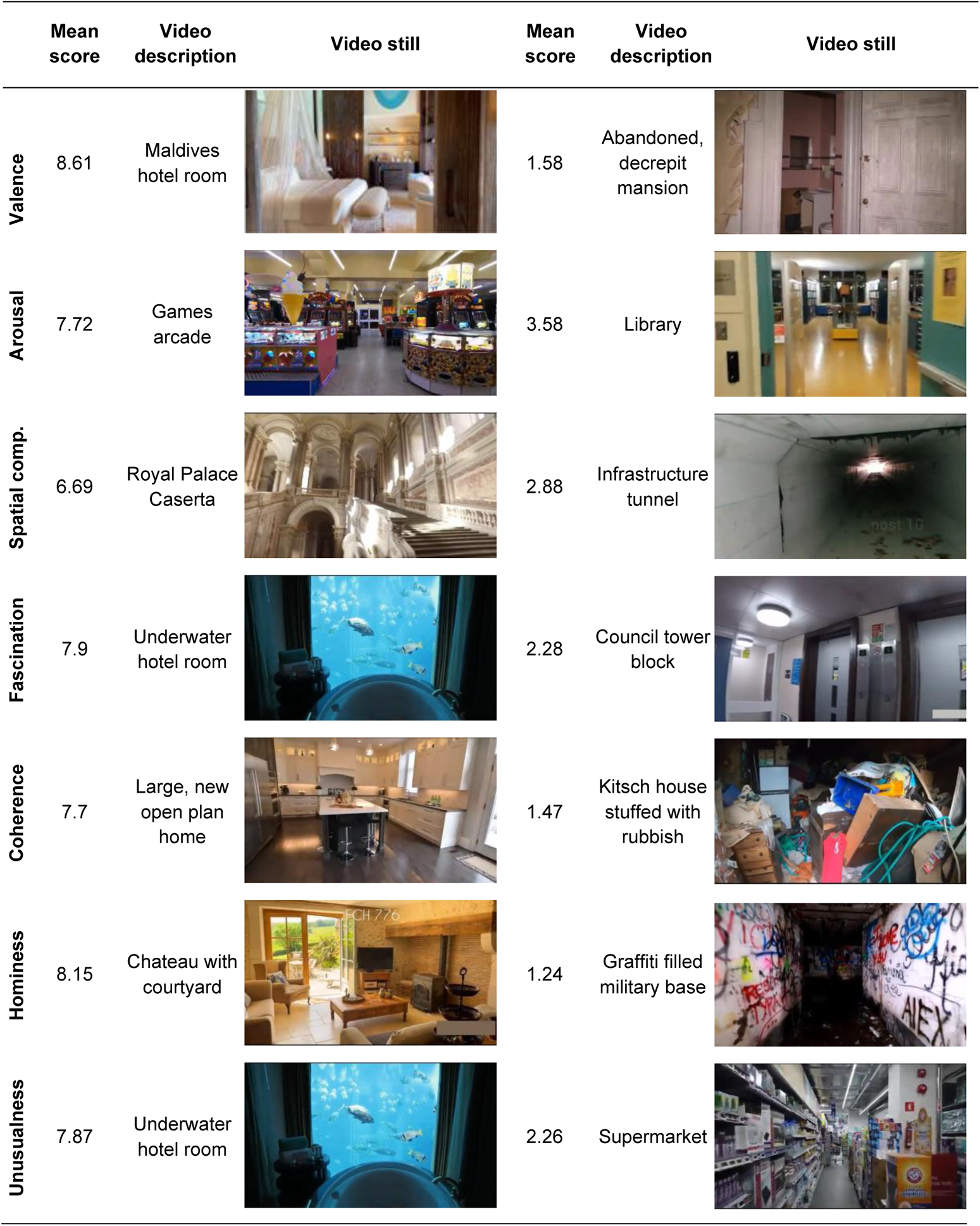
Still images from the highest and lowest mean scoring videos across the seven rating scales.

Examining the distribution of ratings across all participants and all videos, we found that ratings were relatively well distributed across the 1-9 range. The only exception was hominess. Hominess was strongly skewed to low scores, such that some of the videos stood out as being homely, but most were not. Fascination, coherence, spatial complexity and unusualness appeared well spread but had few scores in the middle (rating of 5), so that videos formed two groups, e.g. fascinating places and non-fascinating places, albeit with a good level of variation within these two groupings.

### 3.2. Correlations between ratings

Taking mean scores for each of the 72 videos, we examined the correlation (r) between the seven response measures, along with the presence of green or blue space (calculated using a binary code where 1 = presence of green/blue space, 0 otherwise) and people (1 = people appear, 0 otherwise). The correlation matrix is presented in table 5; a scatter plot matrix of the relationships between the seven key measures is presented in figure 5; and the network between the seven key measures is mapped in figure 6.

**Figure 5:**
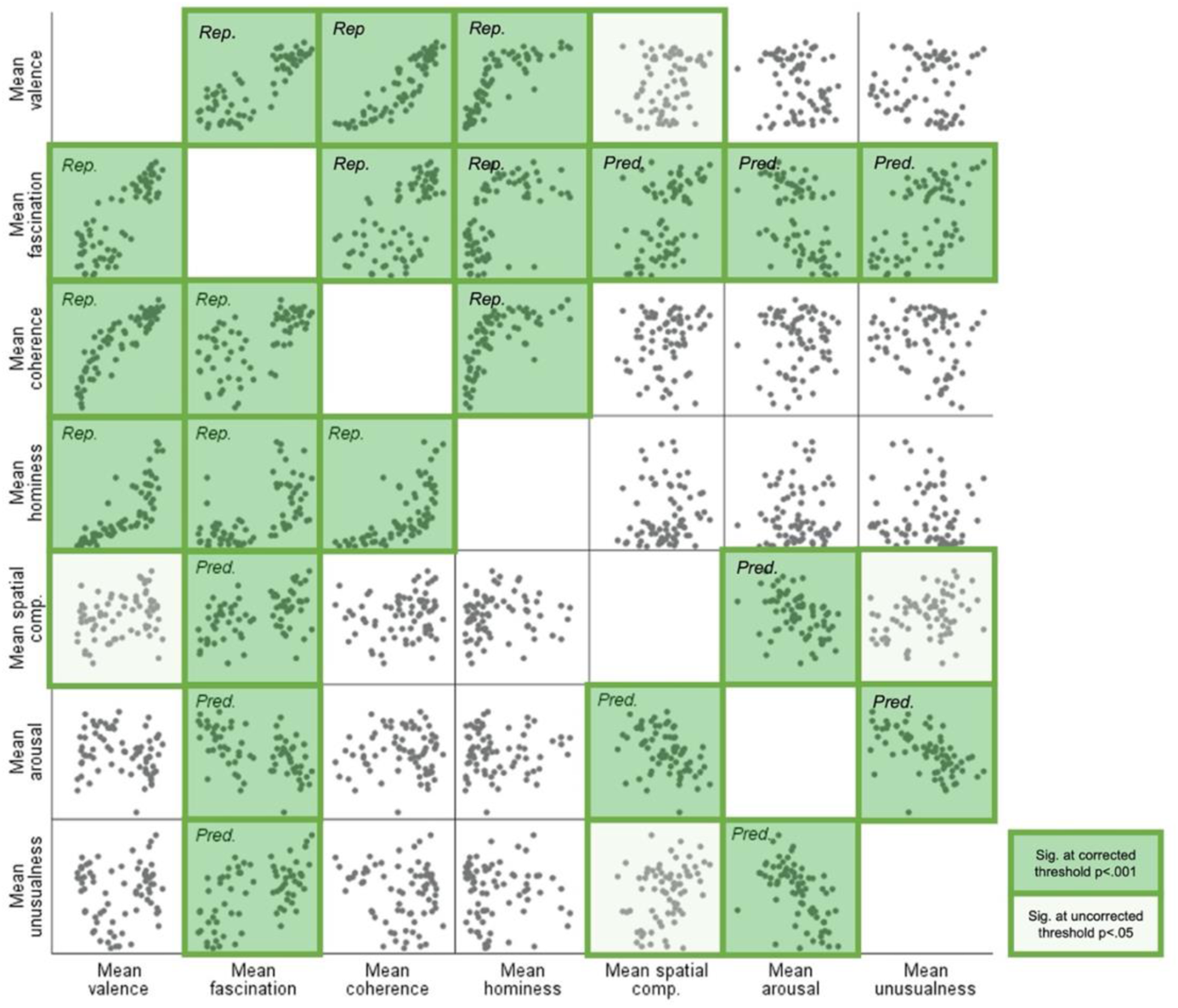
Scatterplot matrix of relationships between each of the seven key measures (N = 72). Rep. = correlations sought to replicate. Pred. = predicted correlations from literature (see table 1). Dark shaded squares indicate significant relationships at the Bonferroni corrected threshold (p < .001). Light shaded squares indicate that correlations between Spatial Complexity and Valence, and Spatial Complexity and Unusualness were significant at p < .05, but these were not predicted and thus did not meet the required threshold for significance.

**Figure 6:**
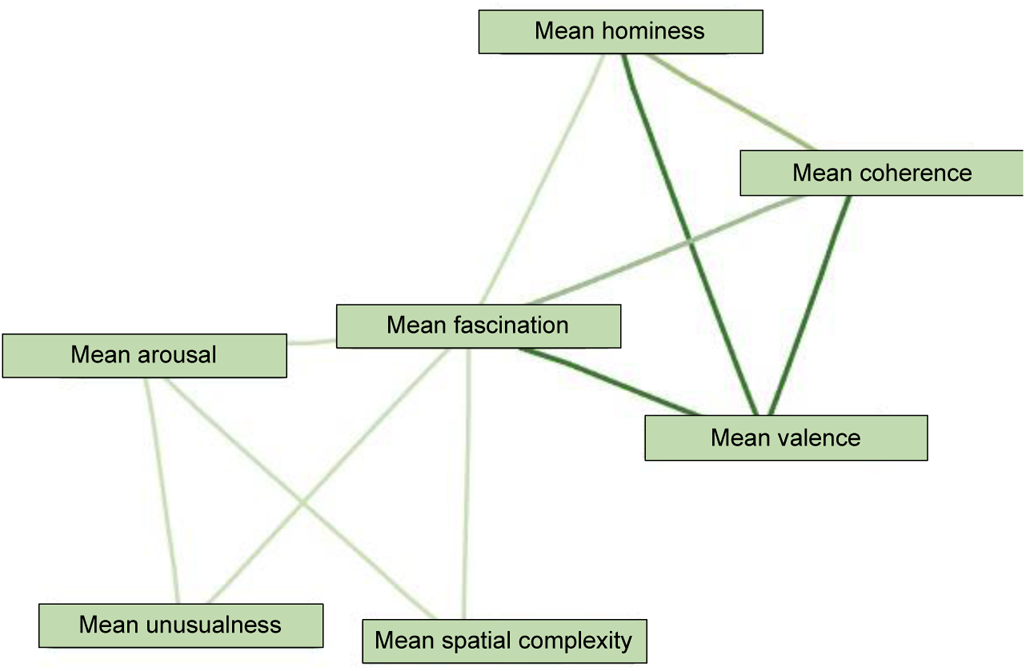
Network map of significant correlations between the primary seven measures. Darker lines indicate stronger correlations. All correlations were positive.

**Table 5:** Correlation Matrix

|  | Mean<br>Valence | Mean<br>Fasc. | Mean<br>Coh. | Mean<br>Hominess | Mean<br>Spatial C. | Mean<br>Arousal | Mean<br>Unusual. | Green/<br>blue | People |
| --- | --- | --- | --- | --- | --- | --- | --- | --- | --- |
| **Mean**<br>**Valence** | 1.00 |  |  |  |  |  |  |  |  |
| *Sig. (2-tailed)* |  |  |  |  |  |  |  |  |  |
| *95% CI* |  |  |  |  |  |  |  |  |  |
| **Mean**<br>**Fascination** | .840** | 1.00 |  |  |  |  |  |  |  |
| *Sig. (2-tailed)* | .000 |  |  |  |  |  |  |  |  |
| *95% CI* | [.755, .897] |  |  |  |  |  |  |  |  |
| **Mean**<br>**Coherence** | .887** | .628** | 1.00 |  |  |  |  |  |  |
| *Sig. (2-tailed)* | .000 | .000 |  |  |  |  |  |  |  |
| *95% CI* | [.825, .928] | [.464, .751] |  |  |  |  |  |  |  |
| **Mean**<br>**Hominess** | .806** | .597** | .681** | 1.00 |  |  |  |  |  |
| *Sig. (2-tailed)* | .000 | .000 | .000 |  |  |  |  |  |  |
| *95% CI* | [.706, .874] | [.424, .728] | [.534, .788] |  |  |  |  |  |  |
| **Mean**<br>**Spatial C.** | .300* | .478** | .160 | .099 | 1.00 |  |  |  |  |
| *Sig. (2-tailed)* | .010 | .000 | .181 | .409 |  |  |  |  |  |
| *95% CI* | [.0734, .4971] | [.277, .639] | [-.744, .378] | [-.136, .323] |  |  |  |  |  |
| **Mean**<br>**Arousal** | .202 | .511** | -.034 | -.020 | .405** | 1.00 |  |  |  |
| *Sig. (2-tailed)* | .088 | .000 | .776 | .868 | .000 |  |  |  |  |
| *95% CI* | [-.0311, .414] | [.317, .664] | [-.264, .199] | [-.251, .213] | [.191, .582] |  |  |  |  |
| **Mean**<br>**Unusualness** | .108 | .564** | -.143 | -.009 | .335* | .564** | 1.00 |  |  |
| *Sig. (2-tailed)* | .368 | .000 | .231 | .940 | .004 | .000 |  |  |  |
| *95% CI* | [-.127, .331] | [.382, .704] | [-.363, .0917] | [-.240, .223] | [.112, .526] | [.382, .704] |  |  |  |
| **Green/blue** | -.021 | -.008 | -.149 | .167 | -.151 | -.096 | .197 | 1.00 |  |
| *Sig. (2-tailed)* | .861 | .947 | .211 | .160 | .207 | .420 | .098 |  |  |
| *95% CI* | [-2.51, .212] | [-.239, .224] | [-.368, .0856] | [-.0673, .384] | [-.370, .0836] | [-.321, .139] | [-.0363, .410] |  |  |
| **People** | .100 | .021 | .200 | -.133 | .036 | -.136 | -.183 | -.290* | 1.00 |
| *Sig. (2-tailed)* | .405 | .863 | .093 | .266 | .762 | .255 | .124 | .013 |  |
| *95% CI* | [-.135, .324] | [-.212, .251] | [-.0332, .413] | [-.354, .102] | [-.197, .266] | [-.356, .0988] | [-.398, .0508] | [-.489, -.0625] |  |

Presence of 'Green/blue' and 'People' in the movies were treated as binary (present/non-present). See table 1 for predictions and statistical thresholds (predicted relationships marked by black boxes). \*\* Double asterisk = relationships meeting criteria for significance ( $p < .001$ ). All predicted correlations were found to be significant. \* Single asterisk = correlations between Spatial Complexity-Valence, Spatial Complexity-Unusualness and People-Green/blue were significant at $p < .05$ , but these were not predicted and thus did not meet the threshold for significance. CI = confidence interval.

The hypothesised three psychological dimensions of architectural experience (fascination, coherence and hominess) have a medium-strong correlation with one another (fascination-coherence r=.628, fascination-hominess r=.597, coherence-hominess r=.681, p<.001). Consistent with predictions, valence is strongly correlated to all three psychological dimensions (valence-fascination r=.840, valence-coherence r= .887, valence-hominess r=.806, p<.001). Below our threshold for significance we also found valence was weakly correlated with spatial complexity (r=.300, p<.05). Interestingly, hominess and coherence correlate with valence and fascination to a similar degree, and also correlate strongly to one another (r=.681, p<.001), suggesting that they might not be easily distinguishable. However, plotting these relationships (Figure 5) hints at their potential nuance - as fascination and valence increase, coherence shows a relatively linear increase, whilst hominess shows exponential increase. Arousal, unusualness, spatial complexity and fascination are all medium-strongly correlated to one another, except for spatial complexity-unusualness which are only weakly correlated and do not meet the threshold for significance (arousal-unusualness r=.564, arousal-spatial complexity r=.405, arousal-fascination r=.511, unusualness-fascination r=.564, spatial complexity-fascination r=.478, p<.001; spatial complexity-unusualness r=.335, p<.05). All of these reported correlations are positive, except between people and green/blue space, which is weakly and negatively correlated (r=-.290, p<.05), indicating that in our videos when green or blue space appears there tend to be less people.

### 3.3. Comparing valence and beauty

As with Survey 1, mean scores were calculated for each rating for each video in Survey 2. The correlation between mean valence scores in Survey 1 and mean beauty scores in Survey 2 is very strong, (r=.937, p<.001), demonstrating that when asked to consider how beautiful or pleasant a space is, results are extremely similar (figure 7).

**Figure 7:**
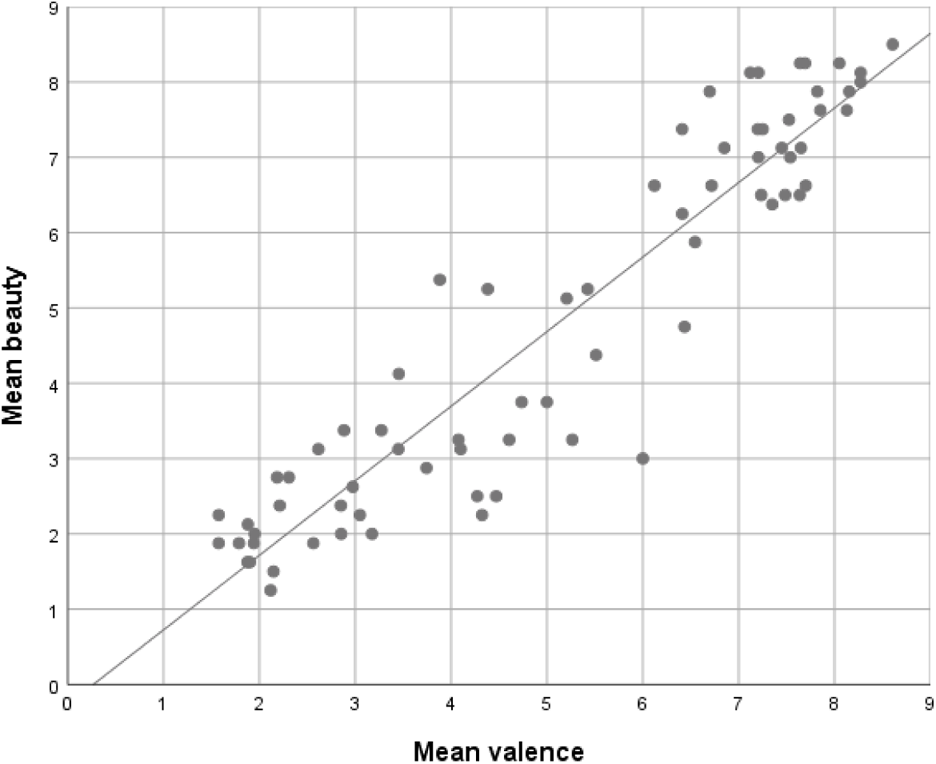
Scatter Plot of mean beauty scores against mean valence scores per video. r = 0.937, p < .001.

For the remaining six qualities, correlations of mean scores between both surveys were largely strong and positive i.e. mean ratings for a given quality in Survey 1 correlate strongly with mean ratings for the same quality in Survey 2 (table 6). The exception to this is arousal (r=.532). For 33 videos, mean arousal scores in Survey 1 were lower than in Survey 2; for 38 videos mean arousal scores in Survey 1 were higher than in Survey 2; and in 1 case they were equal. A histogram demonstrates that mean arousal scores in Survey 1 had a higher concentration around the middle of the rating scale than in Survey 2, where mean scores were more evenly distributed across the scale (figure 8). On further examination, mean beauty and arousal scores in Survey 2 were found to significantly and strongly correlate (r=.828, p<.001), whilst valence and arousal did not correlate in Survey 1 (see table 5).

**Figure 8:**
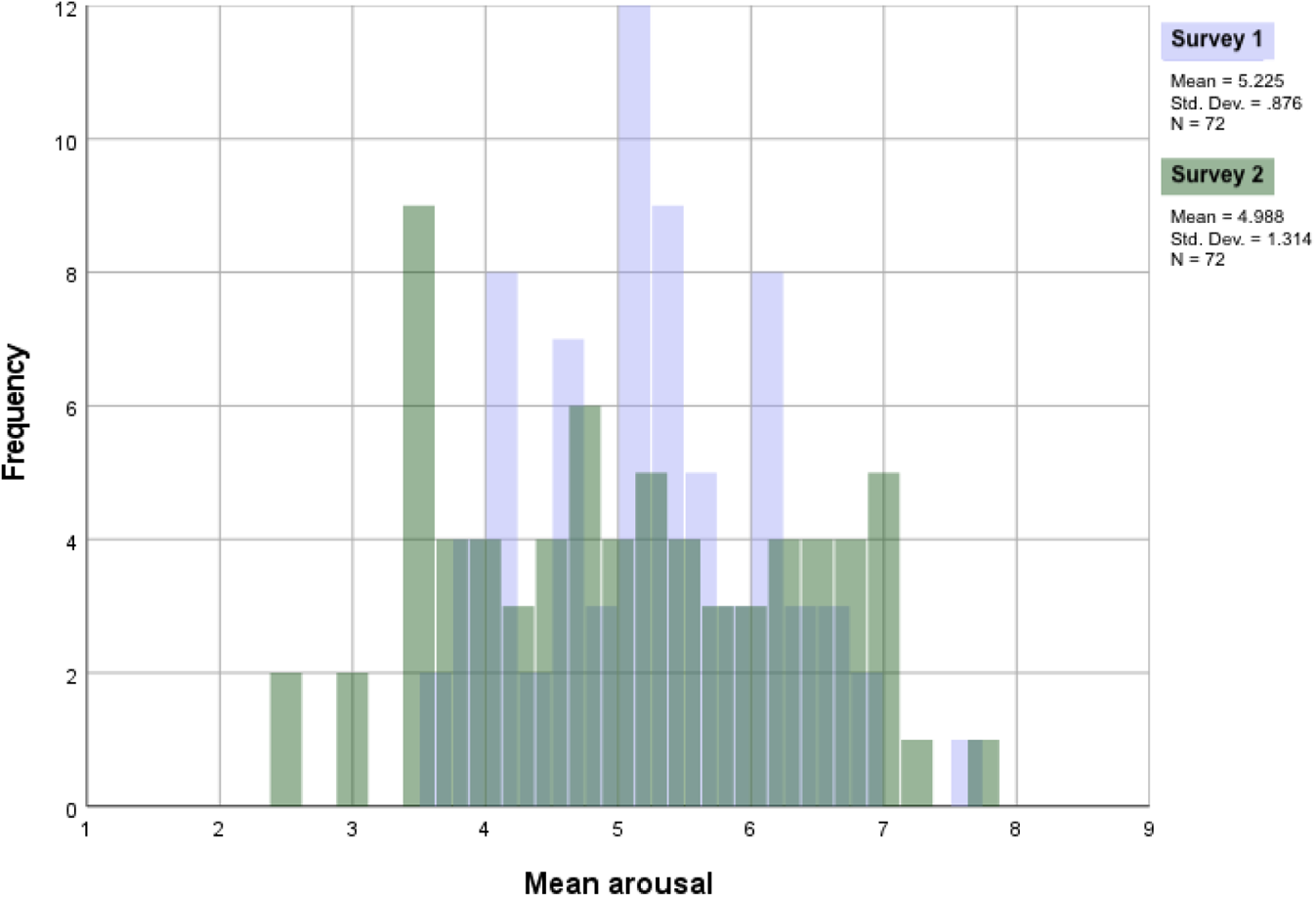
Histogram of mean arousal scores from Surveys 1 and 2.

**Table 6:** Correlations of mean scores of each quality between the two surveys, p<.001

|  | Mean Arousal | Mean Spatial comp. | Mean Fascination | Mean Coherence | Mean Hominess | Mean Unusualness |
| --- | --- | --- | --- | --- | --- | --- |
| Mean of same quality in alternative survey | .532 | .772 | .927 | .918 | .926 | .862 |
| *Sig. (2-tailed)* | .000 | .000 | .000 | .000 | .000 | .000 |
| *95% confidence interval* | [.343, .680] | [.658, .851] | [.886, .954] | [.872, .948] | [.884, .953] | [.788, .912] |

### 3.4. High vs low Navigational Strategy Questionnaire scorers

All participants of Survey 1 answered 12 survey-based strategy questions. Following Zhong (2020), 11 of the 12 questions were averaged to produce a mean Navigational Strategy Questionnaire (NSQ) score indicating ‘competence in cognitive mapping of routes and large-scale environments, and the formation of survey knowledge based on allocentric or environment-centred frames of reference’ (Zhong & Kozhevnikov, 2016). As a metric derived from self-assessment, NSQ scores do not indicate if participants are objectively good or bad navigators, but rather can be used to describe participants’ confidence in mapping and navigating spaces. Table 7 outlines descriptive statistics for NSQ scores across all 73 participants from Survey 1.

**Table 7:** NSQ score descriptives for Survey 1 participants

| Mean | Median | Min | Max | IQ1 | IQ3 |
| --- | --- | --- | --- | --- | --- |
| 2.75 | 2.73 | 1.27 | 4.45 | 2.36 | 3.18 |
| # participants |  |  |  | 18 | 16 |

To examine whether self-perceived navigational abilities influence responses to architectural experiences, data was taken from participants falling in the top and bottom quartiles of NSQ scores. For both groups, mean scores were calculated for each of the seven categories in each video.

Independent 2-tailed t-tests demonstrate that mean scores across all seven categories are not significantly different depending on whether participants have NSQ scores in the top or bottom quartile (p>.05). This suggests that overall, experiences between these two populations are on average similar. On further examination, correlation analyses of mean video scores within these groups helps demonstrate where similarities and differences might lie, according to whether participants believe they have strong survey-based skills or not. See table 8 for correlation matrices in both the top and bottom NSQ-scoring quartiles.

**Table 8:**
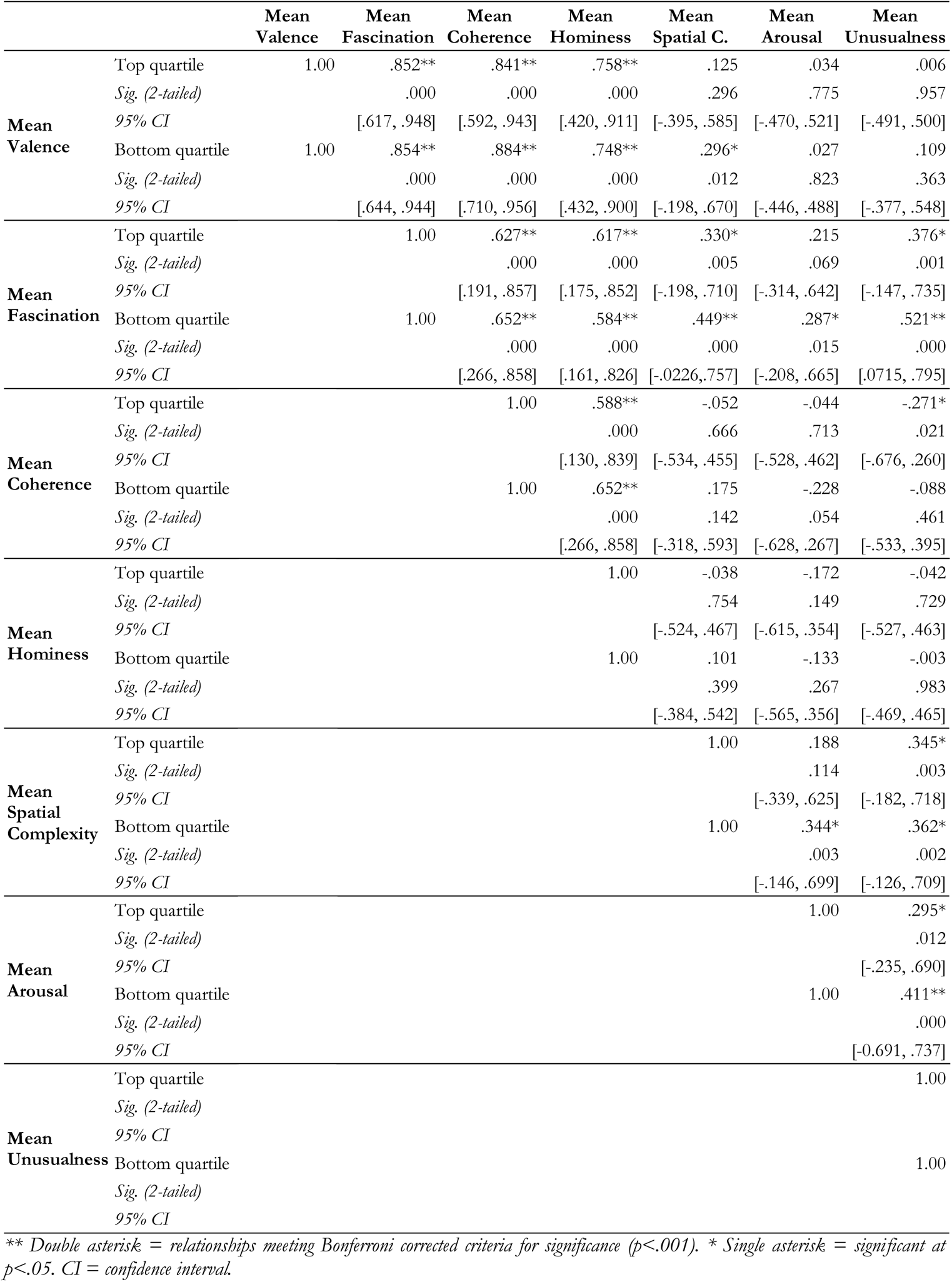
Correlation matrix of mean ratings for top and bottom NSQ quartiles

Between many qualities, relationships are similar whether participants are high or low NSQ scorers. However, there are a number of instances where relationships differ. Arousal only correlates with fascination (r=.287, p<.05) and spatial complexity (r=.344, p<.05) for those in the bottom quartile, and also has a stronger and significant relationship to unusualness amongst this group (r=.411, p<.001). Similarly, spatial complexity correlates with valence, but only for those in the bottom quartile (r=.296, p<.05), and has a stronger and significant relationship to fascination amongst this group too (r=.449, p<.001). Unusualness and fascination correlate stronger and significantly for low NSQ scorers (r.=.521, p<.001). These results support our hypothesis that arousal and spatial complexity are more strongly correlated amongst low NSQ scorers than high NSQ scorers, and further suggests that arousing spaces are more likely to be fascinating and unusual for less confident navigators. For those in the top quartile, coherence and unusualness have a negative relationship (r=-.271, p<.05). We do not find evidence supporting our hypothesis that confident navigators have more positive responses to complex spaces; in fact, we find this to be true for those in the bottom quartile. However these results suggest a sense of spatial comprehension that comes with navigational confidence – whilst both groups see correlation between spatial complexity and unusualness, high NSQ scorers also identify a lack of coherence in more unusual spaces. Overall, these differences in correlations need to be treated cautiously. Many of the relationships that differ between the two groups are not significant at the corrected threshold; they are presented here descriptively in order to outline patterns in the relations for future studies to explore.

### 3.5. Effect of green/blue space and presence of people

Our focus is architectural experience, and thus we wanted to ensure ratings were not overtly influenced by the incidental appearance of green or blue space, or by the presence of people. Thus, two sets of t-tests were conducted. The first evaluated if mean ratings significantly differ between videos with or without green/blue space across all categories (table 9), and secondly if means significantly differ between videos with or without people (table 10).

**Table 9:** Independent 2-tailed t-tests for equality of means (equal variances not assumed) for videos with green/blue space (N=44) vs. videos without green/blue space (N=28)

|  | t | df | Sig. (2-tailed) | Mean difference | Std. Error difference | 95% CI | Cohen's D |
| --- | --- | --- | --- | --- | --- | --- | --- |
| **Mean Arousal** | .758 | 45.4 | .452 | .172 | .227 | [-.285, .629] | 0.194 |
| **Mean Valence** | -.183 | 64.7 | .856 | -.097 | .531 | [-1.16, .964] | -.0438 |
| **Mean Spatial Comp.** | -1.27 | 57.1 | .209 | -.264 | .208 | [-.681, .152] | -.303 |
| **Mean Fascination** | -.067 | 59.0 | .947 | -.029 | .433 | [-.895, .838] | -.0166 |
| **Mean Coherence** | -1.33 | 67.1 | .187 | -.495 | .371 | [-1.24, .246] | -.308 |
| **Mean Hominess** | 1.55 | 69.7 | .126 | .634 | .410 | [-.184, 1.45] | .341 |
| **Mean Unusualness** | 1.66 | 55.6 | .102 | .554 | .334 | [-.114, 1.22] | .403 |

**Table 10:**
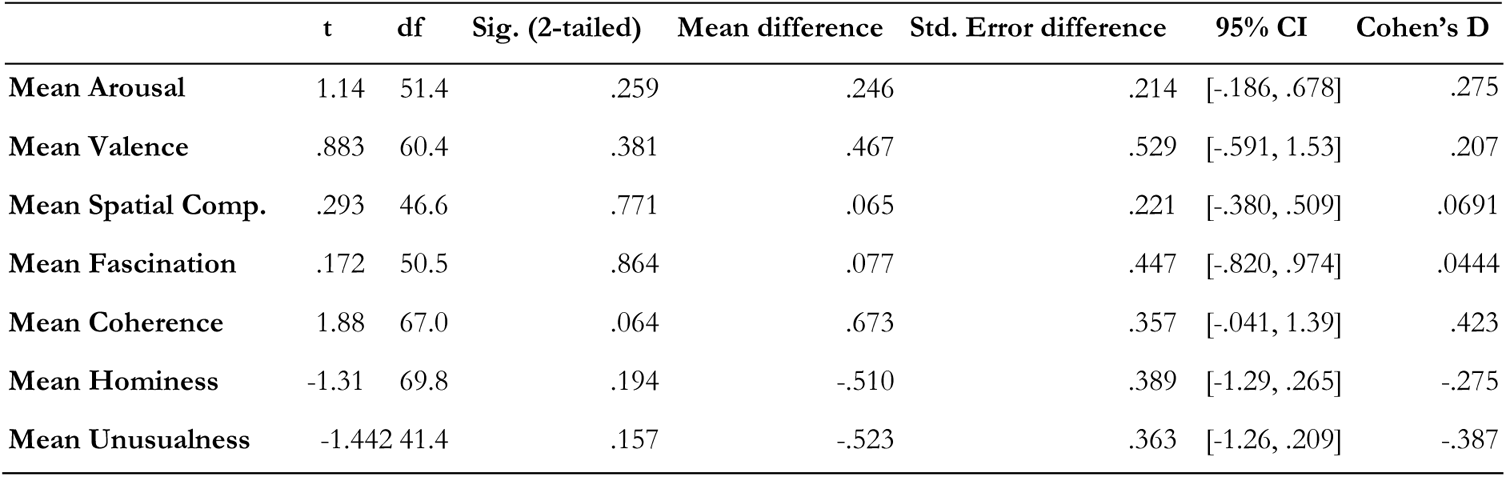
Independent 2-tailed t-tests for equality of means (equal variances not assumed) for videos with presence of people (N=26) vs. videos without presence of people (N=46)

|  | t | df | Sig. (2-tailed) | Mean difference | Std. Error difference | 95% CI | Cohen's D |
| --- | --- | --- | --- | --- | --- | --- | --- |
| Mean Arousal | 1.14 | 51.4 | .259 | .246 | .214 | [-.186, .678] | .275 |
| Mean Valence | .883 | 60.4 | .381 | .467 | .529 | [-.591, 1.53] | .207 |
| Mean Spatial Comp. | .293 | 46.6 | .771 | .065 | .221 | [-.380, .509] | .0691 |
| Mean Fascination | .172 | 50.5 | .864 | .077 | .447 | [-.820, .974] | .0444 |
| Mean Coherence | 1.88 | 67.0 | .064 | .673 | .357 | [-.041, 1.39] | .423 |
| Mean Hominess | -1.31 | 69.8 | .194 | -.510 | .389 | [-1.29, .265] | -.275 |
| Mean Unusualness | -1.442 | 41.4 | .157 | -.523 | .363 | [-1.26, .209] | -.387 |

Means are not significantly different (p>.05) between videos with or without green/blue space across all categories, nor between videos with or without people across all categories (p>.05). It would be beneficial to ensure these relationships are replicable in future work.

### 3.6. Verifying valence

As mentioned in section 1.3, one of our aims was to develop a set of videos that vary in valence but that elicit substantially consistent valence ratings across participants. Towards this end we conducted a two-tiered process.

First, videos were automatically accepted if more than 75% of valence ratings fell in the hypothesised ratings range: 1-4 if negative, 6-9 if positive (note that each video was hypothesised as either positive or negative in valence by the research team). 50 videos fulfilled this requirement and were automatically accepted.

For the remaining 22 videos, the distribution of positive (>5), neutral (=5) and negative (<5) ratings was calculated to assess whether:

a. More than 75% of scores fall in either the neutral-negative (1-5) or the neutral-positive range (5-9).
b. The mean score is significantly different from a mean neutral score of 5 (using a one-sample t-test).

12 videos passed both these tests, and were therefore accepted. On closer inspection, one further video was also accepted as it had a significant t-test result (p<.05) and high percentage of scores in the positive range (73.5%). For 4 of these 13 accepted videos, valence was switched from what was previously hypothesised by the researchers: a gym and arcade (hypothesised to be negative), and a fort in the ocean and dark theatre stairwell (hypothesised to be positive).

Ultimately, 9 videos were rejected (figure 9). For 3 of these, more than 75% of scores did fall in either the neutral-negative or neutral-positive range, however they had non-significant t-test results (p>.05). On closer examination, the percentage of scores falling in an extended neutral range (4-6) were found to be very high (46.2-64.7%), and so coupled with the non-significant t-test result, we could not conclude that they were either positively or negatively valenced. These included a small home with a dilapidated garden, a community centre, and an airport. We had predicted these to be negatively valenced. The remaining 6 videos failed both tests, with less than 75% of scores falling in either the neutral-negative or neutral-positive range, and non-significant t-test results (p>.05). These included a shopping mall, university library, unpreserved temple ruins, bowling alley, museum with ancient artefacts, and ruins in a less well-known part of Pompeii. Though these videos could ultimately not be distinctly identified as negative or positive in valence, all but 2 had over 50% of scores fall in either the positive (>5) or negative (<5) range. In contrast, scores were very evenly distributed for two videos: the university library (38.5% negative, 33.3% positive), and the small home (44.1% negative, 23.5% positive).

**Figure 9:**
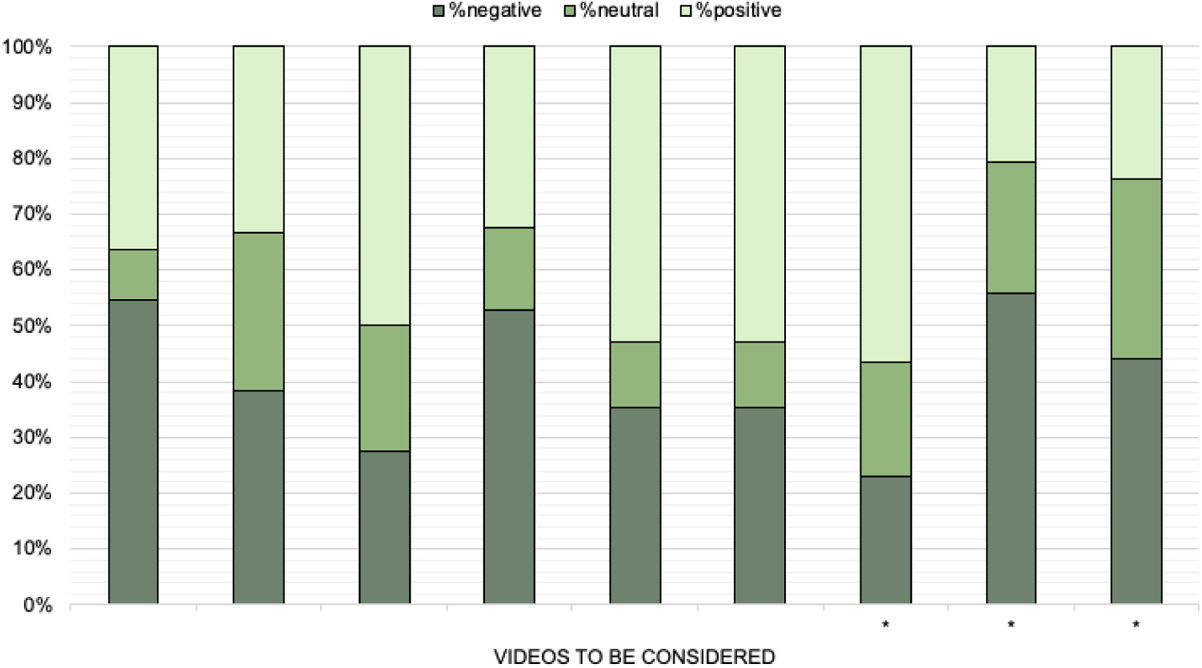
Distribution of negative, neutral and positive scores for the 9 rejected videos. Asterisk indicates videos with >75% of scores falling into the neutral-negative or neutral-positive ranges. All videos failed the one-sample t-test denoting that mean valence scores are not significantly different from a neutral score of 5.

This two-tiered process resulted in a final dataset of 63 valenced videos of trajectories through built environments that can be used as a resource for studying responses to valenced spaces.

## 4. Discussion

### 4.1. Summary of results

In this study we set out to create and validate a novel dataset of valenced first-person-view videos of trajectories through built environments, and to use this dataset to clarify the relationship of core affect and architectural experience. We followed a multistage process to develop a dataset of 72 videos for our experiment. Through an online survey, participants evaluated the videos along seven rating scales. Correlations revealed that valence, fascination, coherence and hominess all strongly relate to one another, as previously demonstrated by Coburn and colleagues (2020). Results also suggest that arousal, fascination, unusualness and spatial complexity are connected, as we hypothesised from previous connections (e.g. Berlyne, 1970; Mendes et al., 2007; Brown et al., 2020; Ruotolo, Sbordone, van der Ham, 2021). A second survey confirmed that beauty and valence ratings are highly correlated, but that their relationships to arousal may differ. Because past evidence suggests poor navigators may treat aspects of the environment such as landmarks differently (Ishikawa & Nakamura, 2012), we examined whether ratings were affected by one’s self-rated ability to map spaces. Exploratory findings suggest slight differences in regard to how confident and unconfident navigators relate factors, but results need further corroboration. Neither the appearance of green/blue space nor the presence of people significantly affected ratings. Finally, to validate and refine the sought-for dataset of videos, we conducted a two-tiered verification process, which resulted in the rejection of 9 videos, and in a final dataset of 63 valenced videos. This dataset provides a new real-world stimuli set, alternative to the image databases previously used in other studies, and offers new insights into architectural experience.

### 4.2. Core affect and dimensions of architectural experience

Our analysis (section 3.1) yielded a number of relationships between core affect and the three central dimensions of architectural experience. As previously hypothesised, valence was positively and strongly correlated to all three dimensions: fascination, coherence and hominess. Coherence and hominess demonstrated similarities that may have suggested they are considered interchangeable, but on closer examination it was evident that hominess is a highly skewed property for our videos while coherence is expressed over a full range. Overall, our findings support Coburn and colleagues’ (2020) theory of architectural experience, showing that results previously obtained using still images extend to dynamic videos of trajectories through built environments. We also help elucidate the nature of their relationship. Moreover, our supportive findings confirm the validity of the terms fascination, coherence and hominess, which were originally coined by Coburn and colleagues (2020) as labels grouping a series of different aesthetic measures (e.g. interest, comfort, beauty). In their original work, participants were not asked to explicitly rate fascination or coherence, but rather scores for these labels were computed through ratings for sub-measure (note that one of the sub-measures within the category of hominess was a rating on hominess itself, whilst the terms fascination and coherence were inspired by the Kaplans’, 1989, work on landscape aesthetics). Our findings support the view that these are effective descriptors that explain the psychological dimensions Coburn and colleagues (2020) aimed to outline.

Here we also examined the relationship between arousal and the three aforementioned psychological dimensions, to clarify their link to core affect. Our analysis revealed that arousal has a moderate correlation with fascination, but not with coherence or hominess. Thus, while positive valence is consistently correlated with all three dimensions, arousal seems to be tied more specifically to fascination. With our arousal scale going from exciting to sleepy, it seems homey or coherent places can be equally exciting or sleepy, whereas fascinating places tend to be considered more exciting. This finding helps us clarify the nature of fascination and to distinguish it from the other two dimensions of architectural experience when it comes to core affect.

When we consider arousal and fascination together with spatial complexity and unusualness, we found an interesting pattern which supports our hypothesised relationships. Arousal shows a moderate positive correlation not only to fascination but also to spatial complexity and unusualness. Complex spaces, as well as unusual spaces, tend to result in more arousing experiences. Fascination also shows a moderate positive correlation with spatial complexity and unusualness. Thus, complex spaces and unusual spaces will tend to result in more fascinating experiences. The relationship between spatial complexity and unusualness is weaker and does not meet the corrected significance threshold, meaning that both simple and complex places may be considered unusual. Thus unusualness and spatial complexity appear to be two semi-independent factors linked to arousing and fascinating architectural experiences.

This understanding of how complexity and fascination interrelate follows a line of research that can be traced back to the seminal work by Kaplan and Kaplan (1989). They highlight the importance of complexity in landscapes in the context of empirical studies of aesthetic preference, and defend the view that complexity drives engagement with the landscape. Here, by using dynamic videos, we extend their line of research from the complexity of a static landscape image to the complexity of a layout; our results suggest that arousal, or activation, is the affective dimension embedded within complexity driving engagement, and resulting in an experience of fascination. Previous research also indicates that arousal impacts spatial memory and navigational strategies, with increasing arousal creating preference for local cues over global cues (Gardony et al., 2011). We may ask: are more arousing spaces found to be more complex as peripheral information on geometry and layout are prioritised less than local cues? Further study is needed to consider the directionality of this relationship and to decipher how focus shifts between local and global landmarks in our videos.

By considering the pattern of correlations, measures can be loosely grouped into two networks: a valence-linked network, which is akin to previous findings (Coburn et al., 2020), and an arousal-linked network, which encompasses unusualness, fascination and spatial complexity (figure 10). These two networks are united in their strong connection to fascination, but could be considered as somewhat independent. Further analysis is needed to deduce the independence and validity of our arousal network. The separation of valence and spatial complexity may also relate to Ranganath & Ritchey’s (2012) identification of two distinct brain networks that aid human memory: the anterior temporal system which is more object-oriented encompassing brain regions linked to affect, and the posterior medial system which is more strongly linked to spatial memory and spatial processing. It may be possible to test this proposal using our video database with brain imaging. Beyond this our videos may provide a useful tool to explore spatial coding of rich real-world environments that go beyond the attempts to build virtual environments that mimic real places (Ekstrom, Spiers, Bohbot, Rosenbaum, 2018; Brunec et al., 2017). Past neuroimaging research has shown the utility of using videos of real-world spaces to simulate navigation and detect brain activity linked to complexity (Howard et al., 2014; Javadi et al., 2017). A limitation of these past studies was that they did not systematically control for the temporal dynamics of valence (e.g. entering a clear or filthy street), which our new video set would afford for future studies.

**Figure 10:**
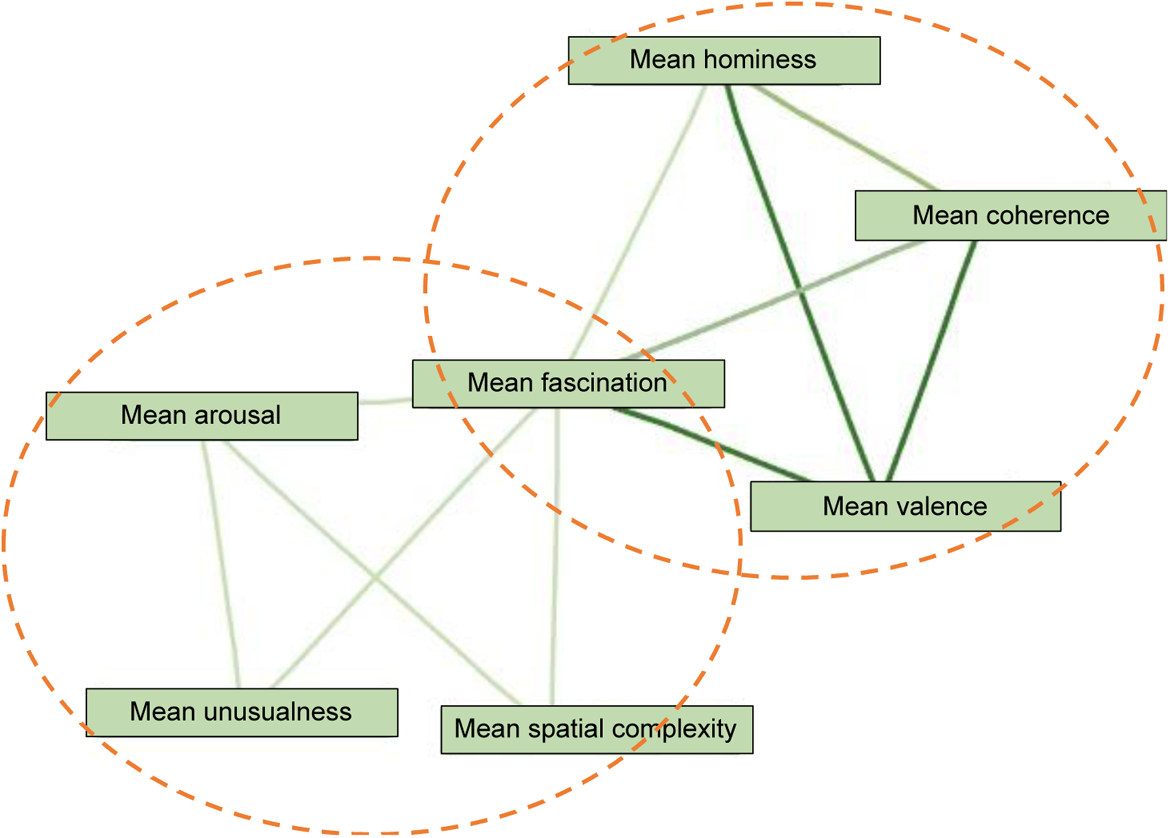
The two overarching groups identified within the primary seven measures. One network connects arousal to other dimensions, and the other network links attributes to valence.

### 4.3. Valence and beauty

Valence and beauty ratings were highly correlated across the two samples compared, suggesting that people consider these two dimensions similarly when it comes to architectural experience. If a video is of a highly positive place it will very likely also be considered a beautiful place. This implies that variables linked to positive affect may also reflect processing of beauty. Our findings follow previous work on varied image databases that similarly show beauty and pleasure ratings to be highly correlated (e.g. Vartanian et al., 2013; Brielmann & Pelli, 2017; Brielmann & Pelli, 2019). Previous studies suggest that this relationship stands regardless of stimulus exposure duration, which aligns with our video stimuli ranging from 31-40 seconds in length (Brielmann, Vale & Pelli, 2017; Belfi et al., 2019). However, despite growing empirical evidence that beauty is a kind of pleasure, they cannot be considered equivalents; e.g. Brielmann & Pelli (2019) find that over 40% of variance in beauty ratings cannot be explained by pleasure ratings. This is important to maintain as our dataset is used in future work.

Examination of the videos revealed that the weakest correlation between mean valence and beauty ratings was derived from videos involving an opera theatre lobby in very dark lighting, a community centre gym, and a gaming arcade. We can hypothesise that this discrepancy could be attributed to semantic distinctions in space types; semantic information has been found to be highly influential in establishing preference. Real-world scenes evoke semantic information describing context and meaning, and when this information is interpreted similarly across a group, there tends to be greater agreement on preference judgments (Vessel & Rubin, 2010). For example, an arcade may be pleasant, but not beautiful, whilst a dark theatre may be beautiful, but unpleasant to be in due to the atmosphere created by the conditions. This is somewhat at odds with Brielmann & Pelli’s (2019) results indicating that whilst unpleasant images can be beautiful, very pleasant images are always deemed beautiful. We could hypothesise that this difference comes from the format of engagement too; contemplating images in front of oneself is a different experience to projecting oneself into a space, where small details - a nice chair or favoured material - might have more prominence and increase pleasantness, but not translate to overarching beauty.

We found that mean ratings for fascination, coherence, hominess, spatial complexity and unusualness correlated very strongly between the two surveys, but that mean ratings for arousal had a weaker correlation. This could be attributed to the difference in how arousal ratings were probed – in Survey 1 as part of the affect grid (Russell et al., 1989), and in Survey 2 as a 1-9 independent rating. One of the benefits of the affect grid is that the two affective dimensions of valence and arousal are given context and meaning, allowing participants to better understand what an increase in arousal may manifest as in relation to a range of everyday words (e.g. ‘sleepy’). However this additional context was lacking in Survey 2, as there is no framework outlining the relationship of beauty and arousal. Therefore it could be argued that mean arousal scores differed between the surveys as the contextualisation of the affect grid in Survey 1 allowed for ratings to be more nuanced and subtler, leading to a larger portion of arousal scores falling in the middle region of the scale. Alternatively, this difference could be attributed to the prior question itself – whether considering valence or beauty. As mean arousal scores in the beauty survey tended to have greater spread than in the valence survey, this could indicate that spaces elicited broader arousal responses in consideration of their beauty than in consideration of their pleasantness. To further investigate, a third survey would be needed, where valence and arousal are rated as independent 1-9 ratings.

In our study valence and arousal did not significantly correlate, whilst beauty and arousal showed a strong positive relationship. This aligns with the valence-arousal model of affect, which considers pleasure and arousal to be independent (Posner et al., 2005). Whilst Brielmann & Pelli (2019) found beauty ratings to be unrelated to arousal ratings across their varied image dataset, on examination of scene-based images only, they noted a positive linear relationship, but of weaker strength than our results suggest. This may indicate that arousal is important in distinguishing how beauty and valence differ specifically in evaluation of scene-based contexts.

### 4.4. High vs low Navigational Strategy scorers

In a more exploratory analysis, we compared scores given by the most confident and least confident navigators in our population. T-tests revealed that across all seven categories, mean video scores calculated by participants falling in the lower quartile of NSQ scorers were not significantly different from mean video scores calculated by participants falling in the upper quartile of NSQ scorers. However correlations suggested some differences in these two group experiences. Our first hypothesis that more confident navigators would have more positive responses to complex spaces is not supported; in fact, less confident navigators appear to have more positive responses to complex spaces. However we find that in high NSQ scorers, coherence negatively correlates with unusualness. Whilst both high and low NSQ scorers find complex spaces more unusual, the more confident navigators are able to identify a lack of coherence in these unusual spaces. This may suggest that confident navigators are better able to make sense of more complex spaces, though interestingly we find no relation between spatial complexity and coherence directly. Our results also potentially support our second hypothesis - that poor navigators may find spatial complexity more arousing. For this group, arousal and spatial complexity also connect more strongly to fascination and unusualness, suggesting that the arousal network identified in figure 10 is perhaps heightened in less confident navigators. However, these results need corroboration: we cannot claim directions of causality, and given that some relationships are not significant at the corrected threshold, these exploratory results need to be taken with caution.

### 4.5. Validating a novel dataset of valenced videos

To develop a dataset of valenced videos of trajectories through real-life built environments, we conducted a multi-step validation process, considering each video’s score spread, and whether mean scores were significantly different from a mean neutral score of 5. Ultimately, 9 videos were rejected, making for a final dataset of 63 valenced videos.

Our results from the selection process may give some insight into what environments lead to mixed responses. The 9 rejected videos included a small home with a dilapidated garden, a community centre, airport, shopping mall, university library, unpreserved temple ruins, bowling alley, museum with ancient artefacts, and ruins in a less famous part of Pompeii. Many of these spaces are not conventionally beautiful or awe-inspiring environments. However, many do have semantic associations (e.g. enjoying travel, seeing friends). As discussed before, semantic information is extremely important in preference making (Vessel & Rubin, 2010). The inconclusiveness or weaker ratings given to these environments could be a product of positive or negative semantic information coupling with weaker aesthetic impact. This hypothesis could even extend to our more surprising rejections - that of the ruins and museum. Though these place types may be stereotyped as more revered due to their historical importance, it is possible that our participant pool reflected those that, for example, do not particularly care about museums or artefacts. Moreover, these places were presented to participants without explicitly indicating their locations; perhaps these spaces would be considered differently with context explaining their more ‘rundown’ appearances. Another possibility is that the sequence of movies created the conditions for this pattern. For example, after seeing a well-kept and sunny beachside resort, an unfamiliar broken ruin in Pompeii may not be seen as a positive place. Future research exploring videos in context with the other videos would be useful for exploring this further. Interestingly, only two videos produced extremely mixed responses: the university library and small home. These were perhaps the most unexciting and commonplace spaces, where some would focus on positive associations (e.g. homely cosiness; educational opportunity), and others more negative (e.g. rundown appearance; stressful studying). Gaining more in-depth, qualitative data on these space types would help identify why they drive mixed responses.

Nonetheless, the resulting set of 63 videos will provide a reliable resource for future empirical studies of architectural experience, confirmed by the aforementioned results replicating findings previously attributed to still images (Coburn et al., 2020). As argued in section 1, emotion is central to the study of architectural experience (Mallgrave, 2013; Zisch, Gage and Spiers, 2014), so it was particularly important that the videos in the dataset resulted in consistently positive or negative valence ratings across participants.

### 4.6. Limitations and future directions

The central thesis to this study lies in employing a set of first-person-view videos of trajectories through architectural environments to better our understanding of architectural experience. Video was considered a useful medium as it is as widely accessible as images, similarly presenting scenes that are completely realistic, whilst also incorporating an additional layer of experience – that of movement through a given space. While we believe this work constitutes a useful development in the types of stimuli that can be used for empirical studies of architectural experience, there are nonetheless still some notable differences between our stimuli and real-life architectural experiences. In contrast to images, videos are dynamic, allowing a viewer to experience a space as they would in reality, however the route and pace of our journeys are set by default. The viewer follows a predetermined path, whereas real-world architectural experience is less passive, as a viewer can and must often decide what paths to take through spaces. A related limitation is that videos are solely visual, while real-life experience is distinctly multimodal, involving non-visual senses from audition to proprioception. We added no sound as the videos we sourced often had a commentary. Future work could address this by creating bespoke videos and possibly through the use of 360 degree or 3D film footage with sound presented via immersive, photo-realistic virtual reality (VR) or augmented reality (AR) (Mazumder, Spiers & Ellard, 2020; Krugliak & Clarke, 2022). It would also be possible to extend this to wireless hmdVR to explore immersion and a more natural experience, and to use laser-scanning of environments for porting in VR, then manipulating the architecture in the models (Zisch et al., 2022). This would also allow for study of peripheral vision, which may be key to garnering a sense of atmosphere (Pallasmaa, 2012).

Our ratings capture snapshot responses to the overall experience of each built environment. The reality of architectural experience is that feelings are likely to ebb and flow along a journey, as no real-world experience is uniform from start to finish. Our ratings do not identify moments within each trajectory where a participant’s feelings might change, however this is something that can be explored in future research through the gathering of continuous ratings. Some of our videos had clear events where the affect increased or decreased. For example, turning to see the room opens to a beautiful ocean view, or opening a door to a room full of cluttered junk. It would be interesting in future work to predict which events individuals would most react to, from gathering sufficient other prior responses.

More fine grain ratings would also be useful for examining brain imaging data recorded during movie-viewing. For example, the DMN appears to track internal state during the course of an ongoing aesthetic experience, increasing in activity over low interest stimuli, thereby suggesting a disengagement and return to rumination and non-stimulus evoked cognition (Belfi et al., 2019). However, it has been found that continuous ratings and singular ratings post-stimuli tend to correspond highly when viewing still art (Belfi et al., 2019), and that while watching videos of landscapes or dance performances, the process of making continuous ratings does not impact overall ratings (Isik & Vessel, 2019). Therefore we can assume that our post-stimuli ratings are valid. Nonetheless, we have provided the foundation for future work that will be able to unpick more dynamic changes of experience.

We aimed to evaluate which videos had consistent valence across people. In empirical aesthetics, however, it tends to be concluded that there is rarely consistency in what is aesthetically pleasing across people, and that personal taste can create large variances in valence ratings between participants (Vessel, Starr & Rubin, 2012; Brielmann & Pelli, 2019; Belfi et al., 2019). Moreover, aesthetic experience can tap into highly personal neural responses concerned with the self, which are not driven by features of the stimuli but by personal experience (Vessel, Starr & Rubin, 2012, 2013). However, evidence also suggests that shared preference may be more identifiable in real-world settings, like those of our stimuli. Vessel & Rubin (2010) found real-world images to have much greater consistency in terms of visual preference than abstract images due to shared semantic interpretations. Similarly, Kaplan & Kaplan (1989) showed that landscape preference could be described through facets such as naturalness and legibility. Shared preference may be weaker in architectural images than in landscape images (Vessel et al., 2014; Weinberger et al., 2021), however those that share preferences for interior designs appear to share preferences for exterior designs, suggesting some notion of mutual experience by style (Vessel et al., 2014). Vessel et al. (2014) theorise that the distinction between landscape and architectural agreement could be due to landscapes tying to evolutionary needs whilst architecture does not, leading to judgments being made on more personal reasonings. However, it is possible that architecture does hold the same behavioural relevance in the sense that successful navigation is pertinent to reaching goals (finding food, being on time for a meeting etc.). Moreover, in the architectural context, empirical research has identified how certain design parameters covary with neural activity (e.g. ceiling height, curvature; Vartanian et al., 2013, 2015), and the study of health and wellbeing in buildings has garnered a detailed understanding of the importance and preference for features such as biophilia and daylight (e.g. Joye, 2007; Cramer & Browning, 2008; Steemers, 2015; Camargo, Artus & Spiers, 2018). This suggests built environments can elicit shared responses. Various steps were taken within our valence verification process to remove videos that showed high variation, but nonetheless it is important to note that a degree of difference will always be found amongst changing participant pools.

Additionally, though calculating NSQ scores and excluding architectural experts gave us some proxy for individual difference, there are many more facets that could be considered, as individual differences can explain navigational performance and spatial processing (e.g. Coutrot et al., 2022; Spiers et al., 2021; Zanchi et al., 2022). It would be beneficial to conduct more research including participants with different personality types and traits to get a more nuanced understanding of experiences, given that individual differences in personality are known to impact affective responses in general (Winter & Kuiper, 1997). It would also be interesting to determine how those with an architectural training react to our stimuli as they have been shown to react differently to spatial scenes than other professionals (Cialone, Tenbrink & Spiers, 2018).

Another challenge with the approach taken here is that labelling emotion is complex. Questions surrounding what specific words mean and how they are measured have long been debated (Berridge, 2018). Here we were interested in the subjective, conscious experience of emotions that an individual has when inhabiting a space. Thus we explored core affect through valence and arousal, as they are deemed the relevant axes when trying to label feelings concisely in a ‘common space’ (Man & Cunningham, 2021). Nonetheless, tools for measuring affect can be perceived and utilised differently by different participants. Barrett (1998) outlines how people may be more valence-focused or arousal-focused, attending more to one of these two dimensions to inform how they would label their affective experience. Highly valence-focused people seem to adopt a more dimensional model of emotion by using affect descriptors of similar valence less distinctly and more concurrently, whilst arousal-focused people use discrete affective descriptors more pointedly. This creates friction as to whether dimensional or discrete theories of emotion are most appropriate for self reports of feeling. Barrett (1998) concludes that there is no singular answer, but that it may be misleading to look at discrete emotional categories (e.g. sadness, fear, anger - all of which are negatively valenced) when it is known that some (valence-focused) people may not report them as separate feelings. Our use of the dimensional affect grid avoids this issue by not asking participants to consider differently labelled emotions, and still benefits from having affective descriptors as guiding anchors around the grid to help shape meaning. Ultimately, no model can capture the breadth of emotional experience, and individual differences will always emerge in terms of how a model is used and self reports expressed. It is also worth noting that self reports of emotion are always entangled with multiple personal qualities and background demographics (e.g. life experience, cultural background, present mood etc.). Thus, it will be useful to explore the extent to which these results generalise across different groups and cultures.

Aesthetic processing is also highly tied to movement perception and is known to engage the motor system. This is an important consideration in evaluations of aesthetic experience with our stimuli. As Freedberg & Gallese (2007) outline, people have embodied responses to visual stimuli, whether that means one feels the muscular tension of a contorted figure represented in painting, or simulates the gestural brushstroke actions of the artist. Aesthetic processing and motor activity are also intertwined in viewing abstract artworks (Sbriscia-Fioretti et al., 2013) and landscapes (Di Dio et al., 2016) – our brains reconstruct actions and activate motor representations that relate to the images that are being viewed. With paintings of nature, sensorimotor activation relates to imagined immersion and exploration in the painted scenes (Di Dio et al., 2016). This relationship is particularly important to consider with our video database, where affective responses will relate to both the embodied simulation described in studies looking at still scenic images, and the intrinsic dynamism of the video’s movement through the scene. In future brain imaging studies that use our database of dynamic stimuli, it will be important to consider the rationales behind motor responses.

Further analysis can be done on the videos themselves at different levels and in a dynamic way. At a low level, there is the option of conducting a frame-by-frame analysis of the videos (e.g. Global Image Properties, such as self-similarity, of individual frames in the video and salience, see e.g. Coburn et al., 2020; Yesiltepe et al, 2021). This can be followed up with examination of how these measures change as the video progresses (e.g. changes in self-similarity over time across the frames of a single video). At a more conceptual level, evaluations by architectural experts can be used to determine global properties of the built environments, as well as changes in those properties as the first-person view traverses different areas within a built environment (e.g. changes in the openness of the space). In turn, these findings can be applied to architectural design, providing architects with information on the psychological effects of dynamic changes in architectural elements. Overall, the emergence of this novel dataset of videos will be beneficial both for architectural design and for empirical research in psychology and neuroscience.

## 5. Conclusions

We developed a novel dataset of first-person videos of trajectories through built environments, and employed it to explore the connections between core affect and the three central psychological dimensions of architectural experience: fascination, coherence, and hominess. We found that fascination, coherence and hominess are all strongly correlated with valence, as previous work suggests (Coburn et al., 2020). We found no correlation between arousal and coherence or between arousal and hominess, but we found that arousal is moderately correlated to fascination, as well as to spatial complexity, and to unusualness. Together, these findings further our understanding of the underlying dynamics of architectural experience by clarifying the nature of fascination in distinction to coherence and hominess and in relation to its core affective components. Moreover, these results suggest that parameters central to spatial mapping and navigation (spatial complexity) are embedded in the affective and aesthetic processing of built environments.

Central to the present study was the development of a novel dataset of videos. We undertook a seven-stage development process, followed by testing and a two-tiered verification process. This resulted in a final dataset of 63 affectively charged videos with consistent valence polarity. This dataset provides a proxy for real-world architectural experience that combines the reality of real-world images with the dynamism of virtual environments, offering a novel stimulus set for experimental work on architectural experience.

## Data availability

Demographics, rating data and videos can be found at https://osf.io/kbdvg/?view_only=7001f5a0d12b444f8e3586c5aa897ccc.

## Author contributions

**Lara Gregorians:** Conceptualization, Methodology, Formal analysis, Investigation, Visualisation, Writing - Original draft, Reviewing and Editing. **Pablo Fernández Velasco**: Conceptualization, Investigation, Writing - Original draft, Reviewing and Editing. **Fiona Zisch**: Conceptualization, Investigation, Writing - Reviewing and Editing. **Hugo J. Spiers:** Conceptualization, Methodology, Investigation, Supervision, Writing - Reviewing and Editing.

## Acknowledgements

We thank the Leverhulme Trust Doctoral Training Programme for the Ecological Study of the Brain for funding LG at UCL [grant number DS-2017-026]. We thank the Irish Research Council for funding PFV at TCD [grant number GOIPD/2021/570].

## Appendix

Here are some spaces to use as reference points. Please note, these reference images do not necessarily represent the extreme ends of each scale, but just give an indication of what these qualities mean and what your peers might rate.

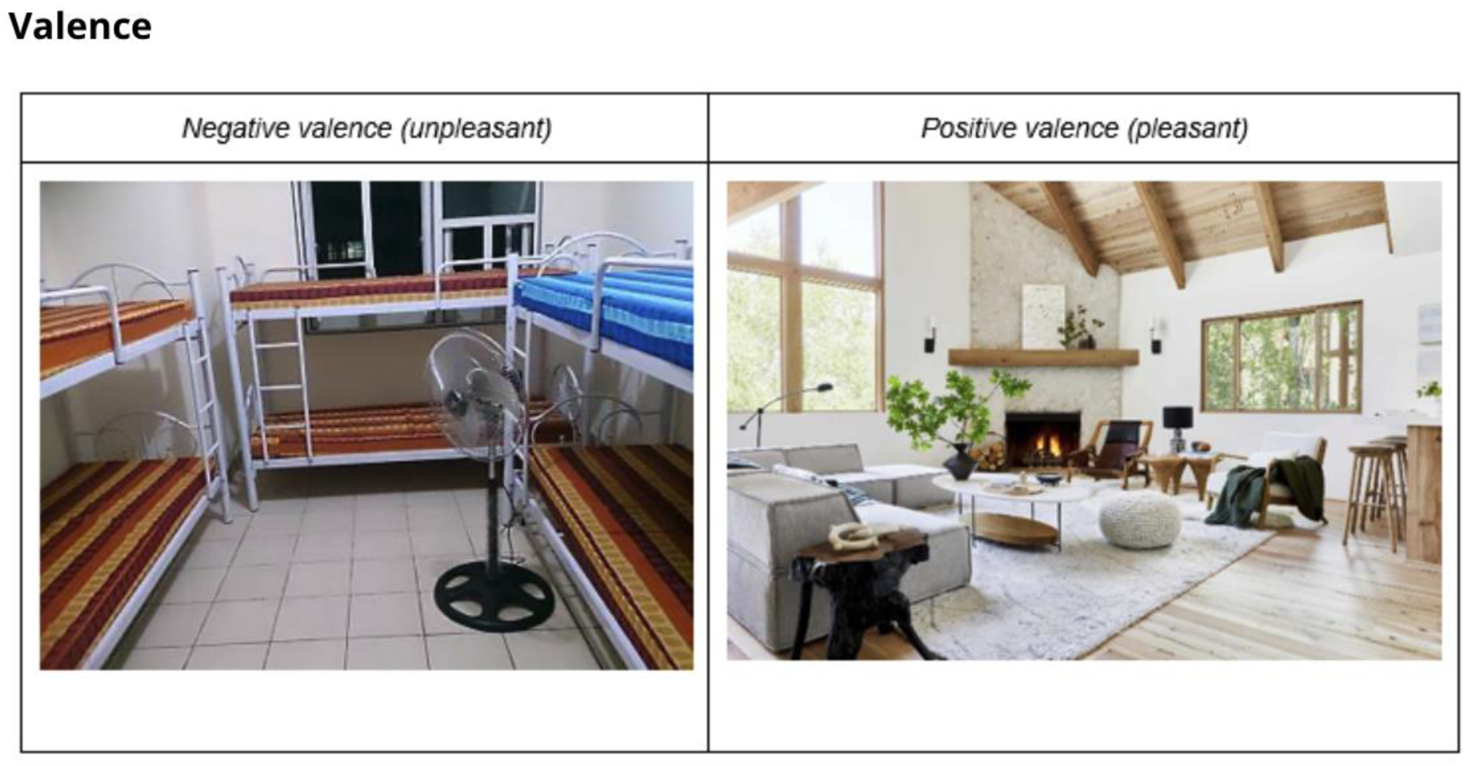

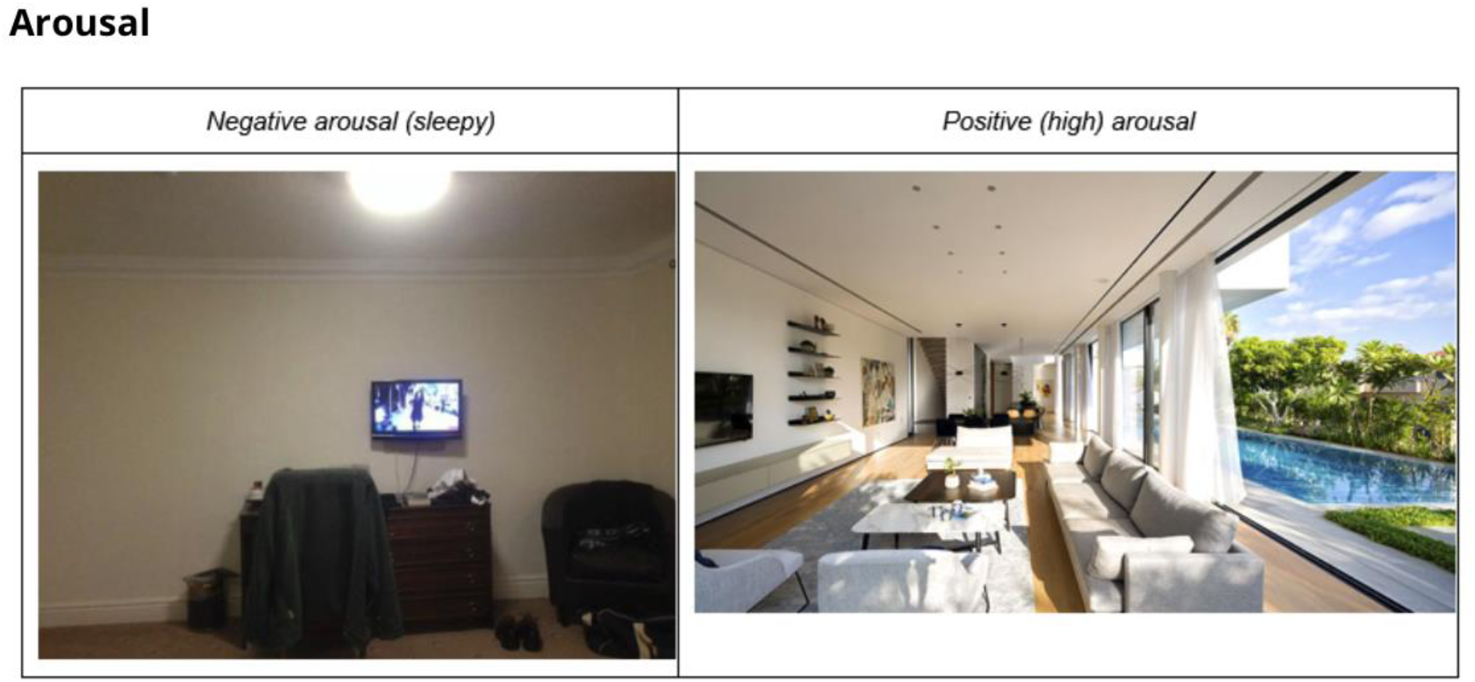

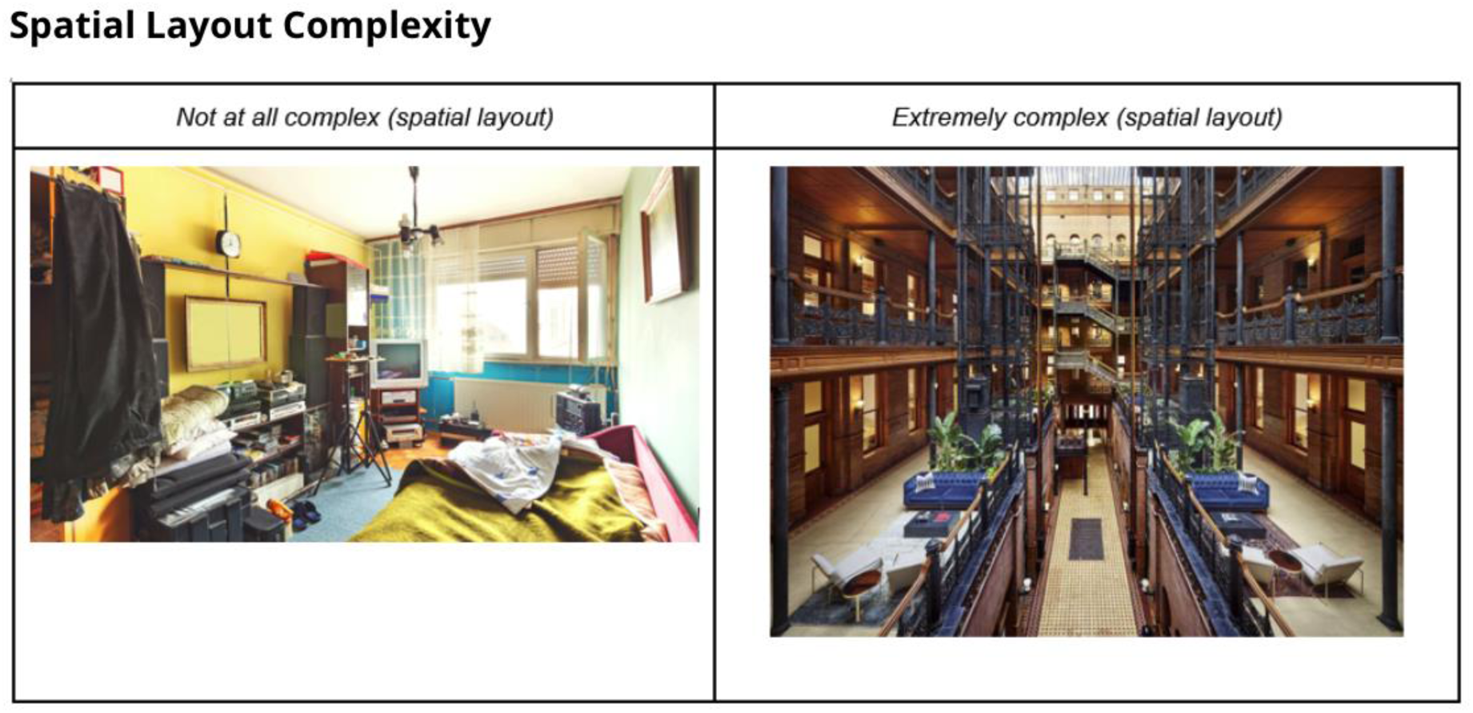

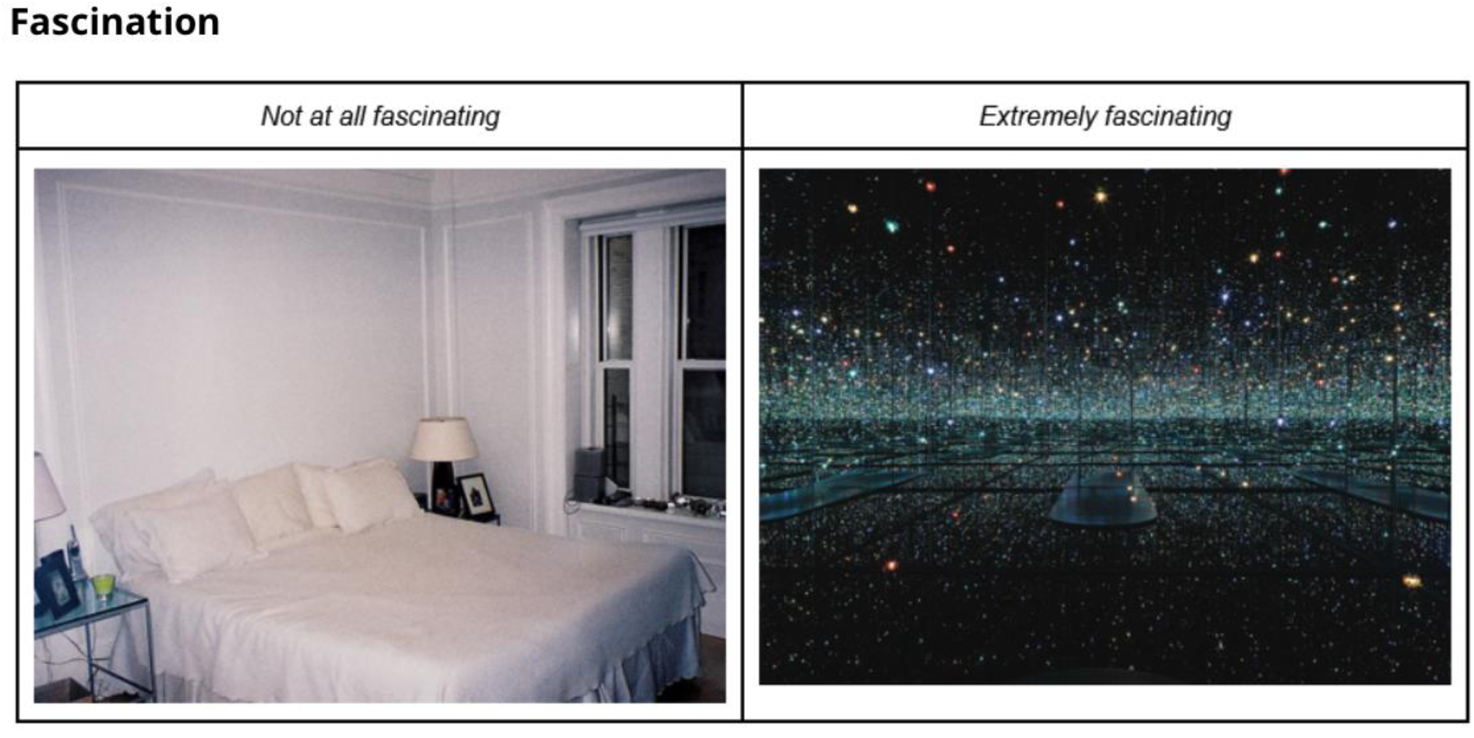

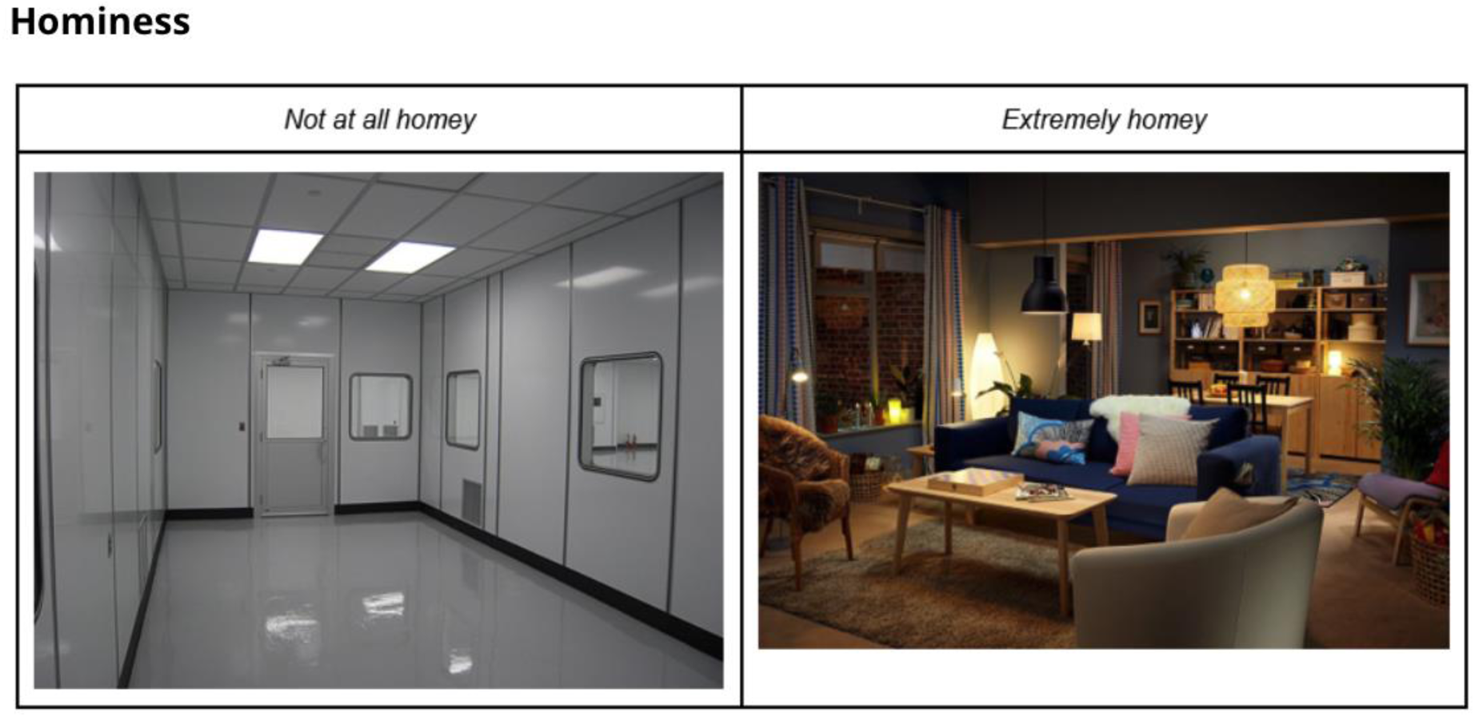

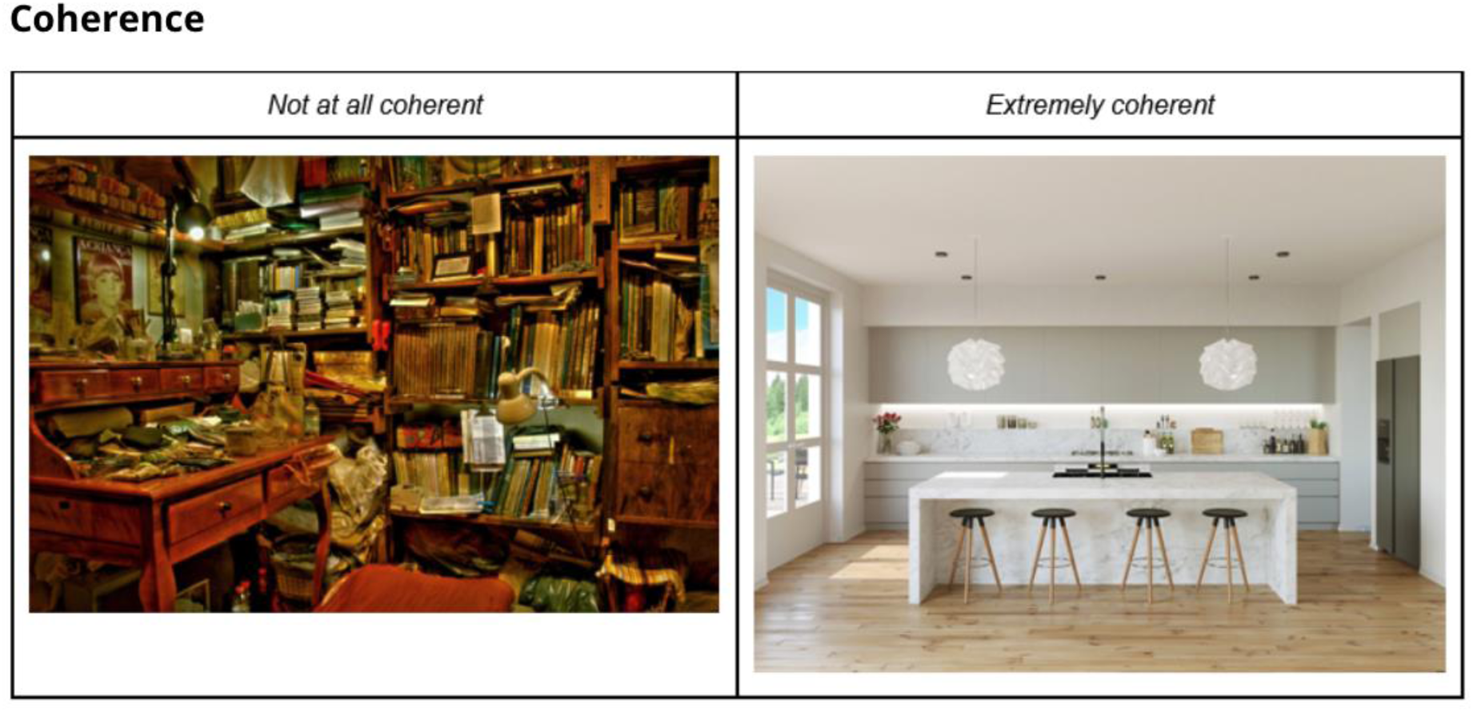

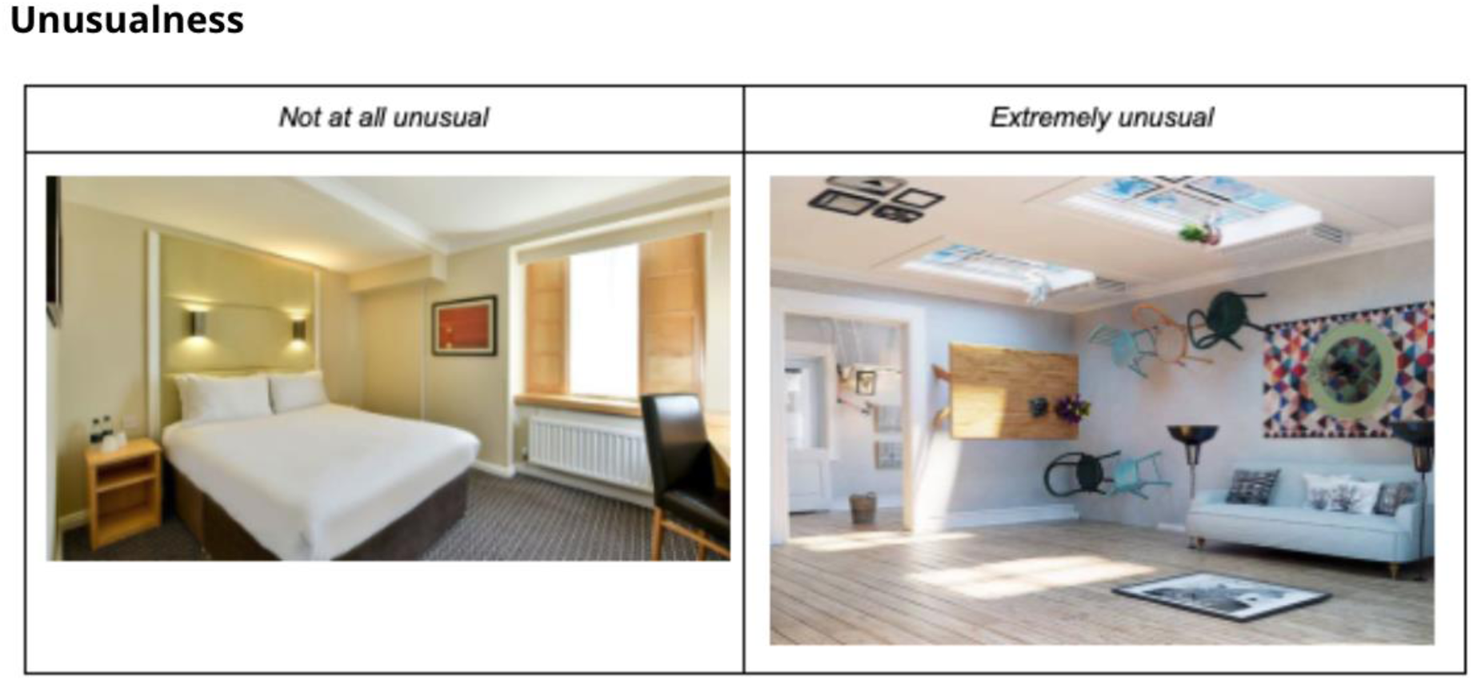

